# Female × male and male × male interactions have limited influence on competitive fertilization in *Drosophila melanogaster*

**DOI:** 10.1101/702720

**Authors:** Stefan Lüpold, Jonathan Bradley Reil, Mollie K. Manier, Valérian Zeender, John M. Belote, Scott Pitnick

**Author notes:** Author contributions: S.L., M.K.M., J.M.B. and S. P., conceived research; S.L., J.B.R., M.K.M., V.Z. and S.P. performed research; S.L. analyzed data; S.L. and S.P. wrote paper. The authors declare no conflict of interest. Data availability: All data will be deposited in the Dryad Repository (doi: XXXXXX).

## Abstract

How males and females contribute to joint reproductive success has been a long-standing question in sexual selection. Under postcopulatory sexual selection (PSS), paternity success is predicted to derive from complex interactions among females engaging in cryptic female choice and males engaging in sperm competition. Such interactions have been identified as potential sources of genetic variation in sexually selected traits but are also expected to inhibit trait diversification. To date, studies of interactions between females and competing males have focused almost exclusively on genotypes and not phenotypic variation in sexually selected traits. Here, we characterize within- and between-sex interactions in *Drosophila melanogaster* using isogenic lines with heritable variation in both male and female traits known to influence competitive fertilization. We found surprisingly few genotypic interaction effects on various stages of PSS such as female remating interval, copulation duration, sperm transfer, or sperm storage. Only the timing of female sperm ejection depended on female × male genotypic interactions. By contrast, several reproductive events, including sperm transfer, female sperm ejection and sperm storage, were explained by two- and three-way interactions among sex-specific phenotypes. We also documented complex interactions between the lengths of competing males’ sperm and the female seminal receptacle, which are known to have experienced rapid female-male co-diversification. Our results highlight the non-independence of sperm competition and cryptic female choice and demonstrate that complex interactions between the sexes do not limit the ability of multivariate systems to respond to directional sexual selection.

**Significance statement:** For species with internal fertilization and female promiscuity, postcopulatory sexual selection (PSS) is believed to depend, in part, on complex interactions between rival males and between the sexes. Although little investigated, clarifying such interactions is critical as they may limit the efficacy of PSS in the diversification of reproductive traits (e.g., ejaculate biochemistry and sperm, genitalia and female reproductive tract morphology). Here, we resolve how sex-specific traits and their interactions contribute to key reproductive events and outcomes related to competitive fertilization success, including traits known to have experienced rapid diversification. Our results provide novel insights into the operation and complexity of PSS and demonstrate that the processes of sperm competition and cryptic female choice are not independent selective forces.

---

Because females of most species mate with multiple males within reproductive cycles (1, 2), sexual selection, encompassing both male-male competition and female choice, can continue after mating in the form of sperm competition and cryptic female choice, respectively (3, 4). As for premating sexual selection, a longstanding goal of studies of post-copulatory sexual selection (PSS) has been to characterize genetic variation in male traits and female preferences that are believed to be the targets of sexual selection, and to identify how such variation relates to differential reproductive success. Explicit demonstration of the means by which phenotypes affect fitness is a prerequisite for resolving the selective processes (5, 6). Compared to premating sexual selection, however (7–9), our knowledge of the causal, mechanistic relationships between sex-specific traits and fitness variation under PSS, and so the contribution of PSS to diversification, is relatively scant (10, 11).

In addition to operational challenges, from observing PSS within the female reproductive tract to discriminating between competing sperm (but see, e.g., 12–14), two inter-related aspects of PSS in particular complicate resolving underlying mechanisms and cloud our understanding of its role in both maintaining genetic variation and driving evolutionary diversification. First, sex-specific mediators of competitive fertilization success tend to be multifarious, potentially including a multitude of genitalic, seminal fluid, sperm and FRT morphological, physiological, neurological and/or biochemical traits, any of which may influence sperm transfer, storage, maturation, motility, longevity, or fertilization (15–19). Second, because sperm competition takes place within the FRT, the competitiveness of ejaculates is likely to depend in large measure on their “compatibility”with the female (17, 20, 21). Any variation in the FRT environment may change the conditions under which sperm compete, and therefore, shift the relative competitive advantage between males (22–25). That such female × male interactions can influence patterns of sperm precedence has been demonstrated in diverse species with both internal (e.g., 22, 26–32) and external fertilization (e.g., 33–37). Further evidence for such interactions comes from studies of conspecific sperm precedence (10, 13, 14, 38). In fact, mounting evidence suggests that competitive fertilization may rarely be independent of female effects (4, 39).

To the extent that numerous, interacting sex-specific traits characterize competitive fertilization systems, PSS is expected to favor the maintenance of genetic variation in reproductive characters and to inhibit strong directional selection on specific traits for three reasons. First, having numerous traits contributing to a fitness outcome can dilute the strength of selection on any single trait. Second, the influence of “genetic compatibility” between males and females on mating or competitive fertilization success, rather than on intrinsic male quality attributes (e.g., sperm number), limits directional sexual selection (40–43). Third, many interacting traits provide the requisite conditions for non-transitive competitive outcomes in the manner of a rock-paper-scissors game (44), which further limits the strength of directional selection (45). Non-transitivity in competitive fertilization success has been experimentally demonstrated for *Drosophila melanogaster* by using fixed-chromosome lines (46–48), and for domestic fowl by using artificial insemination (30). All previous studies that have demonstrated either interactions between the sexes influencing the pattern of sperm precedence or non-transitivity in competitive fertilization success have done so without studying any specific traits (i.e., by using genetically discrete lines or geographic populations), or else have examined a single pair of interacting, sex-specific traits, such as sperm length and female seminal receptacle (SR) length (22) or genetic variation in male sex peptide and its female receptor (47).

In order to advance our understanding of PSS, we set the goals of (a) identifying many of the putative sex-specific targets of PSS (e.g., copulation duration, sperm length, number and *in vivo* swimming velocity, and female remating interval, fecundity, sperm-storage organ morphometry, and sperm storage, ejection and use), (b) quantifying their genetic variation, and (c) determining their contribution to variation in competitive fertilization success within a multivariate framework. To accomplish these goals, we embarked on a three-stage research program using isogenic populations of *D. melanogaster* with sperm that express either green (GFP) or red (RFP) fluorescent protein in their sperm heads (12, 25, 49, 50). The fluorescent tags allow direct visualization of living sperm within the FRT while discriminating between sperm from competing males within twice-mated females, as well as tracking the spatiotemporal fate of both males’ sperm throughout remating and progeny production by females. In the first stage of this program, we held the female genetic background (i.e., isoline) constant and competed males from different isolines in order to resolve male-mediated contributions to competitive fertilization success (49). We demonstrated that longer and slower sperm are better at displacing sperm of a previous male from the female SR (i.e., primary sperm-storage organ), or resisting such displacement by incoming sperm of a subsequent mate (49). In the second stage, we resolved female-mediated contributions by holding the genetic background of all males constant while competing their ejaculates within females from different isolines (25). This study revealed how females can strongly bias sperm storage between males by varying the time between remating and ejecting a mass containing excess second-male and displaced first-male sperm from their bursa copulatrix before initiating oviposition (25). Because in *D. melanogaster* paternity is shared among males in proportion to their sperm remaining in storage and being able to compete for fertilizations (12, 25, 38, 49), female sperm ejection is likely to be a key element of cryptic female choice in this species (25). Here, in the third and final installment, we report on experiments in which the genetic backgrounds of both males and females were systematically varied in order to identify genotypic effects and interactions while resolving the contribution of multivariate traits to male × male, female × male, and female × male × male interactions to variation in competitive fertilization success. Specifically, we mated 6 female and 6 RFP male isogenic lines in all possible combinations and, 2–5 days later, allowed each female to remate with a male from one of 3 GFP isogenic lines. Across these 108 different mating trios in 8 replicates, we then tested how genotypic combinations and interactions between male (e.g., sperm length and number) and female attributes (e.g., remating interval or SR length) influence reproductive events known to affect competitive fertilization (e.g., timing of female post-mating sperm ejection or sperm storage).

## Results

In the first set of analyses, we examined how the different genotypes (6 female, 6 first-male and 3 second-male) and any two- or three-way interactions between them contributed to variation in reproductive parameters (e.g., sperm transferred or stored, timing of female sperm ejection). To this end, we used linear mixed-effects models (LMM) or, where stated, generalized LMM (GLMM) with the four temporally separated blocks as a random factor and accounting for overdispersion. Sample sizes for different traits in these genotypic analyses ranged between *N* = 559 and *N* = 761. We used an information-theoretic approach (51, 52) that accounts for model uncertainty by comparing all candidate models derived from a global model (for details see Material and Methods). To reduce uninformative model complexity, we excluded candidate models that were more complex versions (e.g., one additional parameter) of any model with a lower AIC value (51, 53). All model set tables are provided in the Tables S1–S12.

Female remating interval, ranging between 2 and 5 days, was best predicted by female genotype (ΔAIC_c_ = 11.48) (Table S1). The number of 1^st^-male sperm still residing in the FRT at the time of remating was also primarily explained by the female’s genotype, but with some contribution of the first male (ΔAIC_c_ ≤ 0.45, ΔAIC_c_ to next model = 27.0; Table S2). The number of resident sperm at remating is expected to correlate with the number of sperm transferred by first males and by the number of progeny produced by females between successive matings. Indeed, controlling for the remating interval, the number of progeny produced between the first and second mating was influenced by both female and first-male genotypes (Table S3), but no interactive effects (ΔAIC_c_ ≤ 0.63, ΔAIC_c_ to next model = 89.80).

Both the duration of the second copulation and the number of sperm transferred were explained solely by the 2^nd^ male’s genotype (ΔAIC_c_ > 12.16; Tables S4 and S5), whereas the time to female sperm ejection after remating was explained by the female and 2^nd^-male genotypes and their interaction (ΔAIC_c_ = 4.56; Table S6). Total S_2_, which is the proportional representation of both males’ sperm throughout the female’s sperm-storage organs (i.e., both spermathecae and the SR) after female sperm ejection, was explained best by a GLMM including all three genotypes of a mating trio but no interactions (ΔAIC_c_ = 13.42; Table S7). It is important, however, to distinguish between S_2_ of the total FRT and S_2_ of only those sperm occupying the SR (henceforth the “fertilization set”), as the latter are the primary source of sperm for fertilization in *D. melanogaster* (12, 38). The best model explaining S_2_ of the SR included only the female’s and 2^nd^ male’s genotype (ΔAIC_c_ = 14.19; Table S8).

In a second set of analyses (henceforth “traits analyses”), we examined the interrelationships between the male and female reproductive traits themselves, using model inference based on LMMs or GLMMs that included each represented genotype, the female × male × male genotypic combination, and the temporal blocks as random effects. After selecting the confidence model set, we averaged the coefficients using natural model averaging (51, 54) (for details see Material and Methods).

Our first traits analysis focused on the number of sperm transferred by the second males, which we predicted to depend on female size, copulation duration, the number of 1^st^-male sperm residing within the FRT and interactions (49, 55). The parsimonious confidence set consisted of two models on 2^nd^-male sperm transfer (*N* = 558, including all 108 genotypic combinations: ΔAIC_c_ ≤ 0.64, ΔAIC_c_ to next model = 8.50), represented by strong positive effects of copulation duration (*β* = 0.26 (95% confidence interval: 0.10–0.42)) and the number of resident sperm (*β* = 0.48 (0.32−0.63)), and a weak trend for an interaction between them (*β* = 0.25 (−0.05 to 0.55); Table S9). These results suggest that males transfer more sperm when numerous resident sperm are present in the FRT, through longer copulation. Next, we tested the prediction that the time to female sperm ejection should be influenced by the joint effects of SR length and the differences (2^nd^ – 1^st^ male) in sperm length and number between males (*N* = 529, including all 108 genotypic combinations). Here, only the difference in sperm length had an effect (*β* = 0.20 (0.05–0.36); Table S10). Further examination using only the absolute sperm lengths of both males rather than the difference between them revealed that this effect was driven primarily by the 2^nd^ male’s sperm length (LMM, *N* = 529; 1^st^ male: *β* = − 0.01 (–0.09 to 0.10); 2^nd^ male: *β* = 0.10 (0.03−0.17)). This result suggests that longer 2^nd^-male sperm might prolong the time to female sperm ejection (and thus the sperm displacement phase), which was previously shown to increasingly bias sperm storage toward the second male (25).

We further predicted that the relative numbers of first- and second-male sperm stored by females after sperm ejection (i.e., total S_2_) should be explained by the relative sperm lengths and the numbers of 1^st^-male resident and 2^nd^-male transferred sperm at the end of copulation (49), coupled with the timing of female sperm ejection (25). Using candidate models derived from a GLMM including all interactions between ejection time and the between-male differences in sperm length and numbers, respectively, and with a binomial error distribution and logit link (*N* = 505 across all 108 genotypic combinations) as well as an observation-level random effect to account for overdispersion, total S_2_ increased with both ejection time (*β* = 0.43 (0.27–0.60)) and the difference in sperm numbers (*β* = 0.69 (0.51–0.86)), and was further influenced by an interaction between these two predictors (*β* = 0.48 (0.16–0.79); ΔAIC_c_ = 6.99; Table S11 and Fig. S1). This interaction suggests that by delaying or precipitating ejection, females can amplify or dampen, respectively, the competitive advantage of the second male’s larger ejaculate.

As mentioned previously, it is important to discriminate between total S_2_ throughout the FRT and the fertilization set in the SR. Here, we repeated the above analysis for S_2_ in the SR, but additionally included SR length and female thorax length as predictors, to examine more complex links between sex-specific traits explaining relative sperm numbers in the fertilization set. To limit model complexity, we restricted interactions to two- and three-way interactions and further limited all models to a maximum of 10 parameters (including interactions). Although the full model set contained 1294 different models, this was reduced to only 22 models (Table S12) by removing models that were simply more complex versions of any model with a lower AIC_c_ value (51, 53). The resulting confidence model set (ΔAIC_c_ ≤ 6 (56, 57)) consisted of 7 models, with female thorax length (*β* = 0.35 (0.11−0.59)), SR length (*β* = − 0.56 (−0.79 to −0.32)), the time to sperm ejection (*β* = 0.87 (0.63−1.11)) and difference in the number of sperm between males (*β* = 0.76 (0.51−1.00)) having the most important predictors. The difference in sperm length appeared unimportant as a main effect but contributed to all three interactions whose 95% confidence interval excluded zero after model averaging (Table 1 and S12). For example, together with female thorax length and the difference in sperm number, it formed a three-way interaction meaning that, in a small female, any increasing bias in sperm numbers toward the 2^nd^ male will have a strong effect on S_2_ if the 2^nd^ male has relatively long sperm, but a weaker effect if he has short sperm (Fig. S2). In large females, however, the effect of relative sperm length on S_2_ tends to reverse. Further, the interaction between the difference in sperm length and SR length (Table 1 and S12) means that if second males has shorter sperm than their rivals, any increase in SR length reduces S_2_, whereas SR length has a much weaker effect on the fertilization set if the second male has relatively longer sperm (Fig. S3). This result corroborates Miller and Pitnick’s (22) findings using populations of *D. melanogaster* with experimentally evolved, exaggerated sperm and SR lengths as well as subsequent studies that also predicted significant effects of interactions between sperm lengths of both males and female SR length on the fertilization set (25, 39, 58).

**Table 1:** Model-averaged coefficients of the analysis on the fertilization set (i.e., S_2_ within the female SR) following sperm ejection, including the standardized effects of the difference (2^nd^ – 1^st^male) in sperm length (ΔSL) and in sperm number (ΔSN), the time to female sperm ejection (EJT), female SR length (SRL) and female thorax length (FTL). *N* = 508, including all 108 genotypic combinations. See Table S12 for full details.

| Parameter | Estimate | SE | 95% CL |
| --- | --- | --- | --- |
| (Intercept) | 2.35 | 0.17 | (2.02, 2.68) |
| ΔSN | **0.76** | **0.12** | **(0.51, 1.00)** |
| EJT | **0.87** | **0.12** | **(0.63, 1.11)** |
| SRL | **-0.56** | **0.12** | **(-0.79, -0.32)** |
| FTL | **0.35** | **0.12** | **(0.11, 0.59)** |
| ΔSL × FTL | **-0.55** | **0.24** | **(-1.02, -0.09)** |
| ΔSL | -0.10 | 0.17 | (-0.43, 0.22) |
| ΔSN × EJT | 0.40 | 0.25 | (-0.09, 0.88) |
| ΔSL × SRL | **0.48** | **0.24** | **(0.02, 0.94)** |
| ΔSL × ΔSN × FTL | **-1.12** | **0.47** | **(-2.03, -0.21)** |
| ΔSN × FTL | 0.21 | 0.24 | (-0.26, 0.69) |
| ΔSL × ΔSN | 0.18 | 0.25 | (-0.31, 0.66) |

Finally, we used a piecewise structural equation modeling approach to better visualize both direct and indirect effects of these numerous male and female traits on one another and, in particular, on the fertilization set. This analysis revealed a complex network of interrelationships (Fig. 1). Not surprisingly, relative sperm numbers in the FRT immediately after copulation were a good predictor of relative sperm numbers in the SR after ejection, confirming previous results (25, 49). Interestingly, longer SRs indirectly decreased S_2_ by storing more resident sperm at the time of remating, contrasting with previous studies suggesting that longer SRs are associated with increased S_2_, particularly when the second male has longer sperm (22). However, as shown in the simpler models above, it seems likely that SR length interacts with other reproductive traits, and as a result, can affect S_2_ both positively and negatively depending on the context (note that for simplicity, interactions between traits are omitted in our structural equation models).

**Figure 1:**
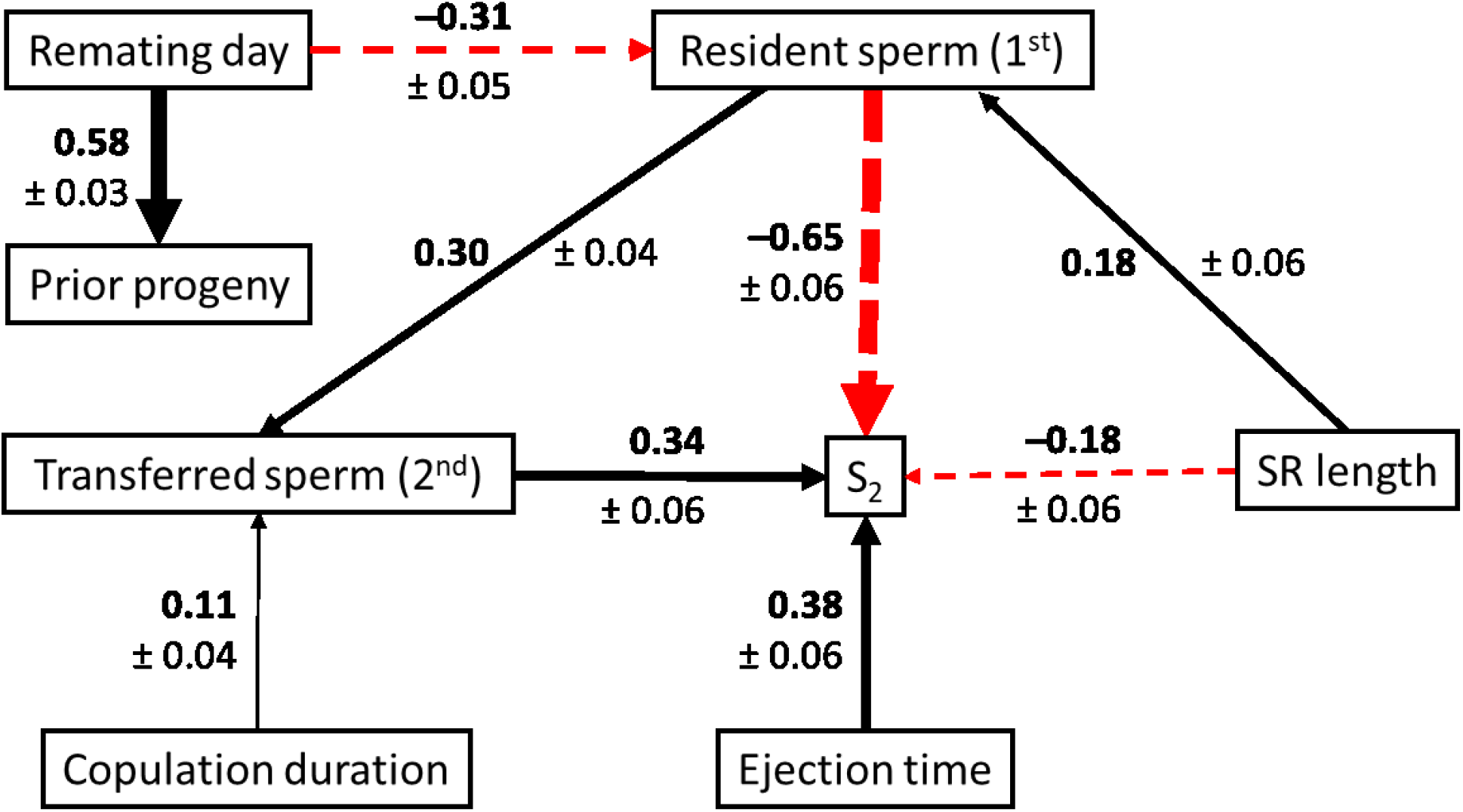
Visual representation of the confirmatory path analysis examining direct and indirect effects on the fertilization set (i.e., S2 within the female seminal receptacle). The width of each arrow is proportional to its corresponding coefficient (in bold, ± standard error). Black, solid arrows depict positive effects, whereas red, dashed arrows represent negative effects. All effects shown were statistically significant (*P* ≤ 0.04); all others (*P* ≥ 0.09) are omitted for better visibility (see Fig. S4 for all relationships examined). This also includes any paths involving the lengths of 1^st^- and 2^nd^-male sperm length and the female thorax, which had significant effects within some of the individual models but not after combining models.

## Discussion

We were able to discriminate which mechanisms of PSS do and do not involve interactions between the sexes, and we resolved some of the sex-specific traits that underlie those mechanisms. It is important to note that these findings are extremely conservative, as with only six respective first-male and female genotypes and three second-male genotypes, we worked with limited genetic variation. Also, numerous sex-specific traits that are expected to contribute to the examined reproductive events were not examined, including genitalic traits, FRT secretions, and male accessory gland proteins (Acps) and their female receptors (18, 21, 59, 60). Hence, although this study represents important progress in understanding the targets and selective dynamics of PSS, there is much more work to be done. It is also important to note that experimental investigation of interactions such as these is complex, and every analytical approach currently available has its own limitations and challenges. Whereas our different statistical approaches resulted in similar patterns, there were quantitative differences in the degree to which female × male interactions explained variation in reproductive outcomes. Because there was some phenotypic variation within the heterozygous F_1_ genotypes for all traits examined, we expect the “traits analyses” to be more sensitive to detecting interactions than the genotypic analyses.

Our investigation suggests that competitive fertilization success in *D. melanogaster* is mediated by both sex-specific mechanisms and by interactions between participants, thus supporting previous studies of this species showing male-specific (e.g., 49, 61, 62), female-specific (25, 63), and interactive effects between the sexes and competing males on reproductive outcomes (e.g., 22, 24, 28, 64). Importantly, the influence of main effects predominated all reproductive outcomes including the fertilization set, which is a consequence of all other traits examined (e.g., the number of sperm transferred by first males, retained and used by females, the number of sperm transferred by second males, and the outcome of sperm displacement activity following remating) and arguably the strongest correlate of variation in male and female fitness (13, 25, 38, 49). As such, our results indicate that ejaculate–female compatibility and non-transitivity in competitive fertilization success might play a more limited role and are unlikely to substantively impede directional sexual selection in this system.

The genotypic analyses suggest, contrary to expectation, that neither the principal fitness outcome (i.e., the fertilization set reflected by S_2_ of the SR) nor the majority of key reproductive events were influenced by genotypic interactions between the sexes. First, the remating interval of females, which is arguably one of the strongest determinants of the intensity of PSS (65, 66), was most plausibly mediated by females alone. This pattern suggests that neither the first male’s transfer of sex peptide and other Acps, which had previously been shown to affect female refractoriness (e.g., 67, 68), nor the second male’s attractiveness (e.g., 69, 70) contributed to variation in the female remating interval. Second, two interrelated traits — the number of progeny produced between successive matings by the female and the number of resident, first-male sperm remaining in storage at the time of remating (which is an important determinant of the fertilization set; Fig. 1) — were both influenced by the female genotype and a weak effect of the first male’s genotype, but not by any interactions. These effects might be attributable to variation in female fertilization efficiency or the timing of sperm ejection (and thus sperm retention) after the initial mating. Further contributing factors might be differences among first males in initial ejaculate size and/or genetic variation in the composition of seminal fluid proteins that are known to influence sperm storage and/or the rate of ovulation and oviposition (e.g., Acp36DE, sex peptide, ovulin (71)). Third, the duration of the second copulation was determined strictly by the genotype of the copulating male, as has been previously shown for *Drosophila* (72, 73). Finally, all three genotypes of a mating trio jointly affected total S_2_, and the more restricted S_2_ of the SR (i.e., the principal fertilization set (12)) was influenced only by the female’s and second male’s genotypes, yet neither variable was influenced by genotypic interactions. We cannot reject the possibility that this lack of interacting effects could be due to limited genetic diversity between the genotypes used in our experiments. However, it is also possible that the compatibility between genotypes themselves generally contributes less to processes underlying competitive fertilization than the interactions between their multiple quantitative traits involved in these processes, as indicated by our ‘traits analyses’.

The only reproductive event with a significant genotypic interaction contributing to its variation was the timing of female ejection. By increasing the time to ejection, females enable more second-male sperm to enter storage and thus further displacement of first-male sperm. Because longer sperm are better at displacing, and resisting displacement by, shorter sperm (14, 22, 25, 58), longer ejection times benefit males with longer sperm and perhaps also the female (39). Here, we found that ejection time was significantly influenced by the female genotype, the second-male genotype, and a female × second-male interaction. Given that sperm ejection is a principal means by which females influence paternity in *Drosophila* (12–14, 25, 74), these collective results suggest that female mediation of competitive fertilization may be more prone to evolving mechanisms entailing genotypic interactions between mating partners than competitive fertilization *per se*.

Results of the traits analyses indicate a more widespread contribution of functional interactions between males and females to variation in reproductive outcomes, including the fertilization set. In this context, it is important to note that previous work with these isolines has demonstrated significant heritability of all of the traits analyzed here, including the number of sperm transferred by males and the timing of female remating and sperm ejection (25, 49). Here we show that, first, the number of sperm transferred by second males — a trait considered to evolve primarily through male-male sperm competition (66, 75) — covaried significantly with the copulation duration and the number of first-male sperm still residing within the female, including a weak interaction between them (Table S9). While male *D. melanogaster* have previously been documented to adjust their ejaculate size to the presence or absence of a competitor’s sperm (55), our results suggest that sperm allocation might be far more sophisticated by responding to the quantity of rival sperm in the FRT, even though the processes underlying such nuanced adjustments remain elusive. Our results further imply that varying numbers of resident sperm, and thus the size of the ejaculate transferred, might explain at least some of the considerable variation in the copulation duration. Second, the difference in sperm length between competing males influenced the timing of female sperm ejection (Table S10), and it further engaged in interactions with female size, SR length, and the difference in sperm number to explain a significant proportion of the variance in the fertilization set (Tables S11 and S12). Although the mechanism underlying the delay in female sperm ejection after mating with a long-sperm male remains unknown, this pattern, combined with a genetic correlation between female sperm ejection and SR length (39), provides a possible functional explanation for the heightened precedence of relatively long sperm in a long compared to short SR reported previously (22) and broadly confirmed here. Our data additionally show that a long SR can also be disadvantageous to the second male by storing more first-male sperm at the time of remating. That greater numbers of sperm may have been occupying the SR at remating might also explain why S_2_ tended to generally lower in long SR compared to short SRs, irrespective of sperm lengths. Thus, these results have potentially far-reaching implications for key elemnts of PSS in *Drosophila*, from male ejaculate investment (see above) to the process of sperm displacement and the interactions of sperm with the female SR or sperm ejection. Despite considerable progress in unraveling the sex-specific contributions to sperm storage and paternity in this model organism, we are a long way from fully understanding the intricacies of this complex interplay.

The present study examined only a small proportion of the sex-specific phenotypes suspected of influencing competitive fertilization success. Nevertheless, all measured male and female reproductive traits contributed to the competitive fertilization set or at least to some reproductive event known to determine it. Further, interactions between competing males and between males and females were shown to explain a significant amount of variation in several key reproductive events, including those generally considered functional components of both sperm competition (e.g., the number of sperm inseminated during remating by a female) and cryptic female choice (e.g., sperm ejection time). Moreover, sperm length, which is known to interact functionally and evolutionarily with SR length (22, 39, 58), did so in a consistent manner in the present study and was further shown to interact with sperm number and SR length in determining competitive fertilization success (i.e., the fertilization set). Our results thus provide further evidence that sperm competition, oftentimes considered to operate between males alone, may in fact rarely be independent of female effects (4, 39). Because the identified interactions included sperm and SR length, which have been shown to represent one of the most extreme co-diversifying systems of male ornamentation and female preference (39), our results demonstrate that multivariate systems with complex interactions between the sexes are not limited in their ability to respond to directional sexual selection. This raises the question of whether the combination of limited non-transitivity between genotypes but a complex interplay between sex-specific, heritable traits might have facilitated the evolution of extreme phenotypes by directional selection while maintaining considerable genetic variation within populations. Finally, our study illustrates both the benefits and empirical challenges of quantifying the contribution of interactions to the operation of sexual selection.

## Materials and Methods

### Experimental material

We performed all experiments with LHm populations of *D. melanogaster* that express a protamine labeled with either green (GFP) or red fluorescent protein (RFP) in sperm heads (12), which permit discriminating sperm from different males and quantifying sperm within the female reproductive tract. Using random individuals from large population cages (all backcrossed to the LHm wild type for 6 generations (12)), we generated isogenic lines (“isolines” (76, 77)) by 15 generations of full-sibling inbreeding, thus yielding theoretical homozygosity levels of 96% (78). To avoid inbreeding effects, we crossed independent pairs of isogenic lines (i.e., males of one isoline with virgin females of another) to create repeatable heterozygous F_1_ genotypes for the experiments described here. Male and female reproductive traits were previously characterized for these crosses and shown to be heritable, and replicated mating trials within given genotype combinations generate repeatable results (25, 39, 49). Based on these assays, we selected parental isolines that captured most of the variance in both male and female reproductive traits among genotypes when creating heterozygotes. In total, our experimental population consisted of six female RFP genotypes, an independent set of six first-male RFP genotypes, and three second-male GFP genotypes. We reared all flies at low densities in replicate vials with standard cornmeal-molasses-agar medium supplemented with yeast, collected them as virgins upon eclosion and aged them for three (males) or four days (females) before their first mating. All males were mated once to a non-experimental female on the day before their first experimental mating to avoid sexual naiveté (24).

### Sperm competition experiment

We have repeatedly shown that paternity (i.e., P_2_) in *D. melanogaster* (including in these isogenic lines) is directly proportional to the respective numbers of sperm from two competing males remaining in storage (S_2_), particularly within the SR, after females have ejected any excess second-male and displaced first-male sperm, thus following a fair raffle among stored sperm (12, 25, 38, 49). Therefore, we focused our efforts on investigating how female × male, male × male, and female × male × male interactions influence the process of sperm displacement until it is interrupted by female sperm ejection (12, 14, 25) and used S_2_ as a proxy of P_2_. Throughout the text, we refer to S_2_ within the SR as the “fertilization set” (*sensu* 79).

Within each of 8 replicates, examined in 4 temporal blocks of 2 full replicate sets (staggered by 2 days), we mated 6 virgin females each to an RFP male and, two days later, to a GFP male in all 108 possible combinations between genotypes (total *N* = 864 trios tested). Females not remating within 4 hours were given additional 4-h remating opportunities on days 3–5 after the first mating. Immediately after the end of the second mating, we removed the male from the mating vial, isolated the female in a glass three-well spot plate beneath a glass coverslip and checked for sperm ejection every 10 min for up to 5 h using a stereomicroscope. We recorded the time to sperm ejection, immediately removed the female from the well and froze it for later quantification of stored sperm, and transferred the ejected mass to phosphate-buffered saline (PBS) on a microscope slide and sealed the coverslip with rubber cement.

For all dissected females, we counted the sperm of both competitors across the different organs of the female reproductive tract (bursa copulatrix, SR, and paired spermathecae with ducts) and determined the total number of sperm for each male in all female sperm-storage organs combined, the proportion of total sperm derived from the first (S_1_) or second male (S_2_), respectively, and the proportion of each male’s total sperm representation in the FRT that reside within the SR. Combining these counts with those of the ejected masses further permitted calculating the number of first-male resident sperm at the time of remating, the number of second-male sperm transferred and both the absolute and relative number of each male’s sperm stored or ejected, respectively.

Finally, for each of the 9 male genotypes, we dissected 6 males after measuring their thorax length, retrieved sperm from their seminal vesicles using a fine probe, fixed the sample on a microscope slide with a mixture of methanol and acetic acid (3/1 v/v), rinsed it with PBS and mounted it under a coverslip in glycerol and PBS (80/20 v/v). We measured the length of 5 sperm per male using the segmented line tool in ImageJ 1.47v at 200× magnification under the dark-field optics of an Olympus BX-60 microscope. For each of the 6 female genotypes, we measured the thorax length of each of 8 females and dissected their reproductive tract into PBS on a microscope slide and covered it with a glass coverslip with clay at the corners, allowing the SR to be flattened to two dimensions without stretching. We then measured SR length using ImageJ at 200× magnification under an Olympus BX-60 microscope with Nomarski DIC optics. Both sperm and SR length are significantly heritable (22, 25, 39, 49).

### Statistical analyses

We performed all analyses using the statistical software package R version 3.4.3 (R Development Core Team 2017). We conducted analyses both at the genotypic and trait level. Due to a lack of specific information on precisely how genotypes or traits should interact to explain focal traits, this study was necessarily somewhat exploratory. Therefore, rather than testing a single model against a null hypothesis, we used an information-theoretic approach to account for model uncertainty (51, 53, 54, 56). Throughout this study, all models were either linear mixed-effects models (LMMs) with the temporal blocking as a four-level random effect or, for the proportional data of S_2_, generalized LMMs (GLMMs) with a binomial error distribution, a logit link, and an additional observation-level random effect to account for overdispersion. Whereas the female, 1^st^- and 2^nd^-male genotypes were the fixed factors (with interactions) in the genotypic analyses, we accounted for genetic non-independence in the traits analyses by including each represented genotype and the female × male × male genotypic combination as random effects.

For each analysis, we generated a model set with all combinations of predictors and interactions (up to a maximum of three-way interactions for interpretability) from a global model using the *dredge* function implemented in the *MuMIn* package (80). We ranked these models by their Akaike Information Criterion with sample size adjustment (AIC_c_ (51, 53, 54, 56)) and limited our confidence model set to candidates within ΔAIC_c_ ≤ 6 of the best model (53, 56, 57), which largely corresponded to cumulative Akaike weights ≥ 0.95. To reduce the retention of overly complex models, we excluded, using the *nested* function in the *MuMIn* package (80), those models that simply represented more complex versions (e.g., one additional parameter) of any model with a lower AIC_c_ value (51, 53).

Although the primary goal was to determine which explanatory variables and interactions were represented in the confidence model set and thus likely to contribute to the variation in the focal trait, we additionally calculated their relative importance as the sum of Akaike weights for the categorical variables of female, first- and second-male genotypes in the genotype-level analyses (51). For terms that are equally represented in the model set, this metric yields higher values for those that occur predominantly in higher-ranked models (51). For the continuous predictors in the traits analyses, we calculated the natural (conditional) averages and 95%confidence intervals of their coefficients as well as their relative variable importance across the confidence model set (51, 54). Here, all predictors were standardized (mean = 0, sd = 0.5) to infer standardized effect sizes (81).

Finally, we used piecewise structural equation modeling (or confirmatory path analyses (82)) in the R package *piecewiseSEM* (83) to visualize how the numerous male and female traits directly or indirectly influence the fertilization set (S_2_ in SR). This approach decomposes a network of relationships into the simple or multiple linear regressions for each response and allows for combinations of different model structures such as LMM and GLMM. Each regression is assessed separately before being combined to evaluate the entire structural equation model (83). The individual models are visualized in Fig. S4.

## Acknowledgments

We thank W.T. Starmer for insightful discussions. This research was supported by a generous gift by Mike and Jane Weeden to Syracuse University and grants from the U.S. National Science Foundation (DEB-1145965 to S.P., S.L., J.M.B., and M.K.M. and DEB-1655840 to S.P.) and the Swiss National Science Foundation (PA00P3-134191 and PP00P3-170669 to S.L.).

## References

1. G. Arnqvist, L. Rowe, Sexual Conflict (Princeton University Press, Princeton, NJ, 2005).

2. M. L. Taylor, T.A. R. Price, N. Wedell, Polyandry in nature: A global analysis. Trends Ecol. Evol. 29, 376–383 (2014).

3. G. A. Parker, Sperm competition and its evolutionary consequences in the insects. Biol. Rev. 45, 526–567 (1970).

4. W.G. Eberhard, Female Control: Sexual Selection by Cryptic Female Choice (Princeton University Press, Princeton, New Jersey, 1996).

5. J. A. Endler, Natural Selection in the Wild (Princeton University Press, Princeton, 1986).

6. E. Sober, The Nature of Selection: Evolutionary Theory in Philosophical Focus (University of Chicago Press, Chicago, 1993).

7. M. Andersson, Sexual Selection (Princeton University Press, Princeton, New Jersey, 1994).

8. M. D. Jennions, A. P. Møller, M. Petrie, Sexually selected traits and adult survival: a meta-analysis. Q. Rev. Biol. 76, 3–36 (2001).

9. H. Kokko, M. Jennions, It takes two to tango. Trends Ecol. Evol. 18, 103–104 (2003).

10. D. J. Howard, S. R. Palumbi, L. M. Birge, M. K. Manier, “Sperm and speciation” in Sperm Biology: An Evolutionary Perspective, eds. T. R. Birkhead, D. J. Hosken, S. Pitnick (Academic Press, San Diego, 2009), pp. 367–403.

11. S. Lüpold, S. Pitnick, Sperm form and function: what do we know about the role of sexual selection? Reproduction 155, R229–R243 (2018).

12. M. K. Manier et al. Resolving mechanisms of competitive fertilization success in *Drosophila melanogaster*. Science 328, 354–357 (2010).

13. MK Manier et al. Rapid diversification of sperm precedence traits and processes among three sibling *Drosophila* species. Evolution 67, 2348–2362 (2013).

14. MK Manier et al., Postcopulatory sexual selection generates speciation phenotypes in *Drosophila*. Curr. Biol. 23, 1853–1862 (2013).

15. R. R. Snook, Sperm in competition: not playing by the numbers. Trends Ecol. Evol. 20, 46–53 (2005).

16. S. Pitnick, D. J. Hosken, T. R. Birkhead, “Sperm morphological diversity” in Sperm Biology: An Evolutionary Perspective, eds. Birkhead TR, Hosken DJ, Pitnick S (Academic Press, San Diego, 2009), pp. 69–149.

17. S. Pitnick, M. F. Wolfner, S. S. Suarez, “Ejaculate-female and sperm-female interactions” in Sperm Biology: An Evolutionary Perspective, eds. Birkhead TR, Hosken DJ, Pitnick S (Academic Press, San Diego), pp. 247–304. (2009).

18. A. Poiani, Complexity of seminal fluid: a review. Behav. Ecol. Sociobiol. 60, 289–310 (2006).

19. I. Carmel, U. Tram, Y. Heifetz, Mating induces developmental changes in the insect female reproductive tract. Curr. Opin. Insect Sci. 13, 106–113 (2016).

20. K. Ravi Ram, M. F. Wolfner, Seminal influences: *Drosophila* Acps and the molecular interplay between males and females during reproduction. Integr. Comp. Biol. 47, 427–445 (2007).

21. L. K. Sirot, M. F. Wolfner, “Who’s zooming who? Seminal fluids and cryptic female choice in Diptera” in Cryptic Female Choice in Arthropods: Patterns, Mechanisms and Prospects, eds. Peretti A V, Aisenberg A (Springer, Cham, Switzerland), pp. 351–384. 1st Ed. (2015).

22. G. T. Miller, S. Pitnick, Sperm-female coevolution in *Drosophila*. Science 298, 1230–1233 (2002).

23. G. T. Miller, S. Pitnick, Functional significance of seminal receptacle length in *Drosophila melanogaster*. J. Evol. Biol. 16, 114–126 (2003).

24. A. Bjork, W. T. Starmer, D. M. Higginson, C. J. Rhodes, S. Pitnick, Complex interactions with females and rival males limit the evolution of sperm offence and defence. Proc. R. Soc. B 274, 1779–1788 (2007).

25. S. Lüpold et al., Female mediation of competitive fertilization success in *Drosophila melanogaster*. Proc. Natl. Acad. Sci. USA 110, 10693–10698 (2013).

26. S. M. Lewis, S. N. Austad, Sources of intraspecific variation in sperm precedence in red flour beetles. Am. Nat. 135, 351–359 (1990).

27. N. Wilson, S. C. Tubman, P. E. Eady, G. W. Robertson, Female genotype affects male success in sperm competition. Proc. R. Soc. B 264, 1491–1495 (1997).

28. A. G. Clark, D. J. Begun, T. Prout, Female × male interactions in *Drosophila* sperm competition. Science 389, 217–220 (1999).

29. T. Nilsson, C. Fricke, G. Arnqvist, The effects of male and female genotype on variance in male fertilization success in the red flour beetle (Tribolium castaneum). Behav. Ecol. Sociobiol. 53, 227–233 (2003).

30. T. R. Birkhead, N. Chaline, J. D. Biggins, T. Burke, T. Pizzari, Nontransitivity of paternity in a bird. Evolution 58, 416–420 (2004).

31. C. Y. Chow, M. F. Wolfner, A. G. Clark, The genetic basis for male × female interactions underlying variation in reproductive phenotypes of *Drosophila*. Genetics 186, 1355–1365 (2010).

32. S. Y. N. Delbare, C. Y. Chow, M. F. Wolfner, A. G. Clark, Roles of female and male denotype in post-mating responses in *Drosophila melanogaster*. J. Hered. 108, 740–753 (2017).

33. E. Turner, R. Montgomerie, Ovarian fluid enhances sperm movement in Arctic charr. J. Fish Biol. 60, 1570–1579 (2002).

34. J. P. Evans, D. J. Marshall, Male-by-female interactions influence fertilization success and mediate the benefits of polyandry in the sea urchin Heliocidaris erythrogramma. Evolution 59, 106–112 (2005).

35. P. Rosengrave, N. J. Gemmell, V. Metcalf, K. McBride, R. Montgomerie, A mechanism for cryptic female choice in chinook salmon. Behav. Ecol. 19, 1179–1185 (2008).

36. L. W. Simmons, J. D. Roberts, M. A. Dziminski, Egg jelly influences sperm motility in the externally fertilizing frog, *Crinia georgiana*. J. Evol. Biol. 22, 225–229 (2009).

37. S. H. Alonzo, K. A. Stiver, S. E. Marsh-Rollo, Ovarian fluid allows directional cryptic female choice despite external fertilization. Nat. Commun. 7, 12452 (2016).

38. M. K. Manier, S. Lüpold, S. Pitnick, W. T. Starmer, An analytical framework for estimating fertilization bias and the fertilization set from multiple sperm-storage organs. Am. Nat. 182, 552–561 (2013).

39. S. Lüpold et al., How sexual selection can drive the evolution of costly sperm ornamentation. Nature 533, 535–538 (2016).

40. T. R. Birkhead, Cryptic female choice: criteria for establishing female sperm choice. Evolution 52, 1212–1218 (1998).

41. S. Pitnick, W. D. Brown, Criteria for demonstrating female sperm choice. Evolution 54, 1052–1056 (2000).

42. T. Tregenza, N. Wedell, Genetic compatibility, mate choice and patterns of parentage: invited review. Mol. Ecol. 9, 1013–1027 (2000).

43. K. P. Oh, A. V. Badyaev, Adaptive genetic complementarity in mate choice coexists with selection for elaborate sexual traits. Proc. R. Soc. B 273, 1913–1919 (2006).

44. J. Maynard Smith, Evolution and the Theory of Games (Cambridge University Press, Cambridge) (1982).

45. A. G. Clark, Sperm competition and the maintenance of polymorphism. Heredity 88, 148–153 (2002).

46. A. G. Clark, E. T. Dermitzakis, A. Civetta, Nontransitivity of sperm precedence in *Drosophila*. Evolution 54, 1030–1035 (2000).

47. R. Zhang, A. G. Clark, A. C. Fiumera, Natural genetic variation in male reproductive genes contributes to nontransitivity of sperm competitive ability in Drosophila melanogaster. Mol. Ecol. 22, 1400–1415 (2013).

48. A. Civetta, A. G. Clark, Correlated effects of sperm competition and postmating female mortality. Proc. Natl. Acad. Sci. 97, 13162–13165 (2000).

49. S. Lüpold et al., How multivariate ejaculate traits determine competitive fertilization success in Drosophila melanogaster. Curr. Biol. 22, 1667–1672 (2012).

50. O. Ala-Honkola et al., Inbreeding reveals mode of past selection on male reproductive characters in Drosophila melanogaster. Ecol. Evol. 3, 2089–2102 (2013).

51. K. P. Burnham, D. R. Anderson, Model Selection and Multi-Model Inference: A Practical Information-Theoretic Approach (Springer, New York)2nd editio. (2002).

52. K. P. Burnham, D. R. Anderson, K. P. Huyvaert, AIC model selection and multimodel inference in behavioral ecology: some background, observations, and comparisons. Behav. Ecol. Sociobiol. 65, 23–35 (2011).

53. S. A. Richards, M. J. Whittingham, P. A. Stephens, Model selection and model averaging in behavioural ecology: The utility of the IT-AIC framework. Behav. Ecol. Sociobiol. 65, 77–89 (2011).

54. C. E. Grueber, S. Nakagawa, R. J. Laws, I. G. Jamieson, Multimodel inference in ecology and evolution: Challenges and solutions. J. Evol. Biol. 24, 699–711 (2011).

55. S. Lüpold, M. K. Manier, O. Ala-Honkola, J. M. Belote, S. Pitnick, Male Drosophila melanogaster adjust ejaculate size based on female mating status, fecundity, and age. Behav. Ecol. 22, 184–191 (2011).

56. M. R. E. Symonds, A. Moussalli, A brief guide to model selection, multimodel inference and model averaging in behavioural ecology using Akaike’s information criterion. Behav. Ecol. Sociobiol. 65, 13–21 (2011).

57. B. M. Bolker et al., Generalized linear mixed models: a practical guide for ecology and evolution. Trends Ecol. Evol. 24, 127–135 (2009).

58. J. M. Pattarini, W. T. Starmer, A. Bjork, S. Pitnick, Mechanisms underlying the sperm quality advantage in *Drosophila melanogaster*. Evolution 60, 2064–2080 (2006).

59. C. E. McDonough, E. Whittington, S. Pitnick, S. Dorus, Proteomics of reproductive systems: Towards a molecular understanding of postmating, prezygotic reproductive barriers. J. Proteomics 135, 26–37 (2016).

60. A. L. Mattei, M. L. Riccio, F. W. Avila, M. F. Wolfner, Integrated 3D view of postmating responses by the *Drosophila melanogaster* female reproductive tract, obtained by micro-computed tomography scanning. Proc. Natl. Acad. Sci. 112, 8475–8480 (2015).

61. A. G. Clark, M. Aguadé, T. Prout, L. G. Harshman, C. H. Langley, Variation in sperm displacement and its association with accessory gland protein loci in *Drosophila melanogaster*. Genetics 139, 189–201 (1995).

62. A. C. Fiumera, B. L. Dumont, A. G. Clark, Sperm competitive ability in *Drosophila melanogaster* associated with variation in male reproductive proteins. Genetics 169, 243–257 (2005).

63. A. G. Clark, D. J. Begun, Female genotypes affect sperm displacement in *Drosophila*. Genetics 149, 1487–1493 (1998).

64. P. D. Mack, B. A. Hammock, D.E. L. Promislow, Sperm competitive ability and genetic relatedness in *Drosophila melanogaster*: Similarity breeds contempt. Evolution 56, 1789–1795 (2002).

65. E. Boorman, G. A. Parker, Sperm (ejaculate) competition in *Drosophila melanogaster*, and the reproductive value of females to males in relation to female age and mating status. Ecol. Entomol. 1, 145–155 (1976).

66. L. W. Simmons, Sperm Competition and its Evolutionary Consequences in Insects (Princeton University Press, Princeton, New Jersey, 2001).

67. T. Chapman et al., The sex peptide of *Drosophila melanogaster*: Female post-mating responses analyzed by using RNA interference. Proc. Natl. Acad. Sci. USA 100, 9923–9928 (2003).

68. J. Morimoto et al., Sex peptide receptor-regulated polyandry modulates the balance of pre-and post-copulatory sexual selection in *Drosophila*. Nat. Commun. 10, 283 (2019).

69. S. Pitnick, Male size influences mate fecundity and remating interval in Drosophila melanogaster. Anim. Behav. 41, 735–745 (1991).

70. D. J. Hosken, M. L. Taylor, K. Hoyle, S. Higgins, N. Wedell, Attractive males have greater success in sperm competition. Curr. Biol. 18, R553–R554 (2008).

71. F. W. Avila, L. K. Sirot, B. A. LaFlamme, C. D. Rubinstein, M. F. Wolfner, Insect seminal fluid proteins: Identification and function. Annu. Rev. Entomol. 56, 21–40 (2011).

72. I. T. MacBean, P. A. Parsons, Directional selection for duration of copulation in *Drosophila melanogaster*. Genetics 56, 233–239 (1967).

73. I. T. MacBean, P. A. Parsons, The genotypic control of the duration of copulation in *Drosophila melanogaster*. Experientia 22, 101–102 (1966).

74. R. R. Snook, D. J. Hosken, Sperm death and dumping in *Drosophila*. Nature 428, 939–941 (2004).

75. G. A. Parker, T. Pizzari, Sperm competition and ejaculate economics. Biol. Rev. 85, 897–934 (2010).

76. P. A. Parsons, S.M. W. Hosgood, Genetic heterogeneity among the founders of laboratory populations of *Drosophila*. I. Scutellar chaetae. Genetica 38, 328–339 (1968).

77. J. R. David et al., Isofemale lines in *Drosophila*: an empirical approach to quantitative trait analysis in natural populations. Heredity 94, 3–12 (2005).

78. D. S. Falconer, Introduction to Quantitative Genetics (John Wiley & Sons, New York). 3rd Ed. (1989).

79. G. A. Parker, Sperm competition games: raffles and roles. Proc. R. Soc. B 242, 120–126 (1990).

80. K. Bartón, MuMIn: multi-model inference. R package, version 1.40.0. Available at: https://cran.r-project.org/web/packages/MuMIn. (2017).

81. A. Gelman, Scaling regression inputs by dividing by two standard deviations. Stat. Med. 27, 2865–2873 (2008).

82. B. Shipley, Confirmatory path analysis in a generalized multilevel context. Ecology 90, 363–368 (2009).

83. J. S. Lefcheck, piecewiseSEM: Piecewise structural equation modelling in R for ecology, evolution, and systematics. Methods Ecol. Evol. 7, 573–579 (2016).

