## Supplementary Tables and Figures for "Female × male and male × male interactions have limited influence on competitive fertilization in *Drosophila melanogaster*"

### Supplementary Figures:

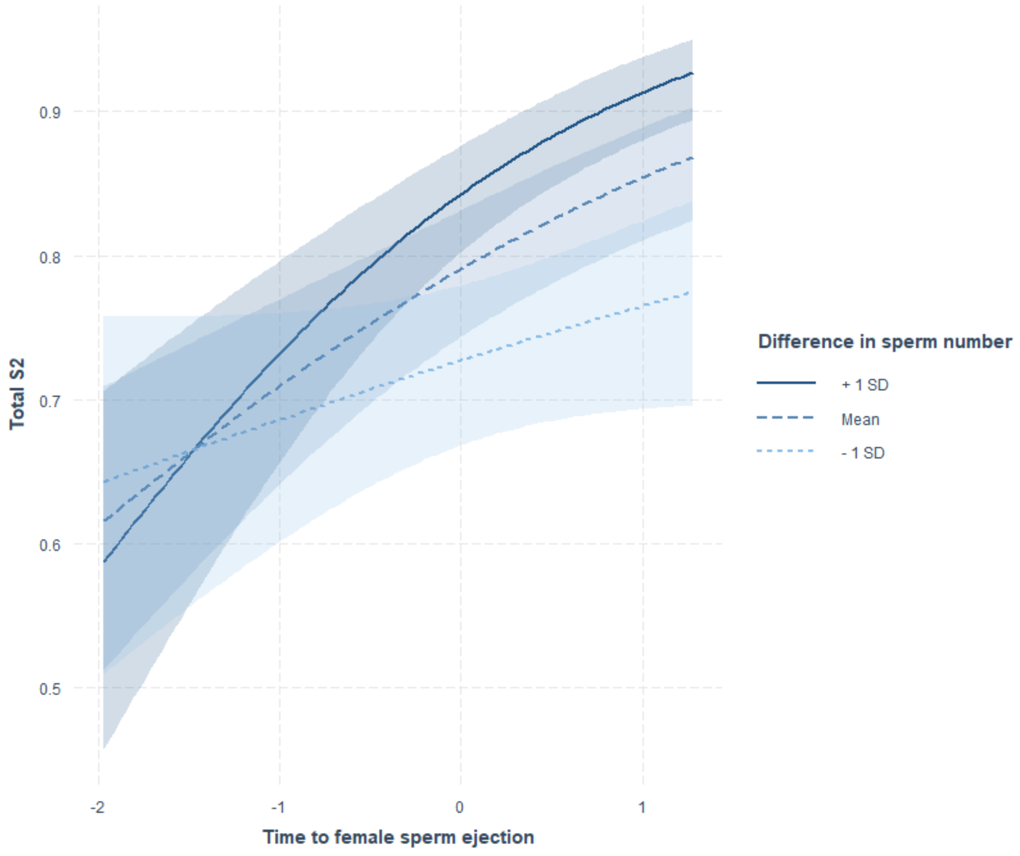


**Supplementary Figure S1:** Conditional effects plot of the two-way interaction between the time from remating to female sperm ejection and the difference (2nd male – 1st male) in sperm number explaining variance in total $S_2$. The plot indicates that the effects of relative sperm numbers and ejection time reinforce one another, with the greatest change in $S_2$ when the second male transfers a disproportionate amount of sperm and the female waits relatively long to eject excess or displaced sperm. The shaded areas around lines depict the 95% confidence intervals.

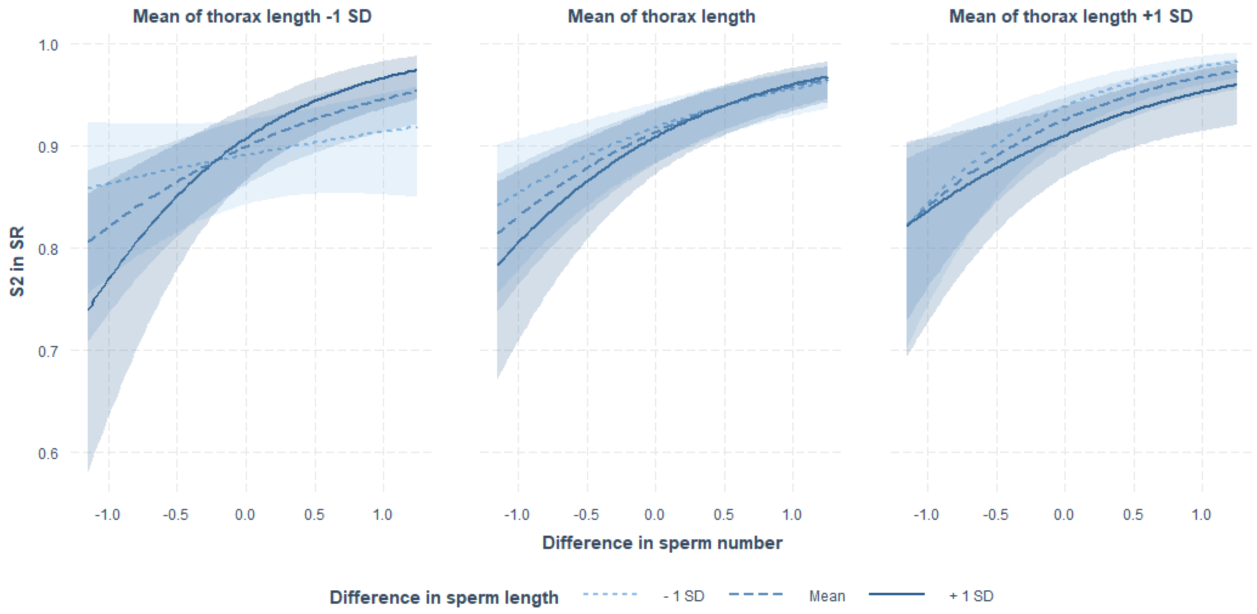


**Supplementary Figure S2:** Conditional effects plot of the three-way interaction between the female thorax length and the relative differences in sperm length and sperm number, respectively, explaining variance in the fertilization set ($S_2$ within the female SR). Differences are shown from the 2nd male's perspective (i.e. 2nd male – 1st male). The plot depicts how any increase in the number of sperm transferred by the second male relative to the first-male sperm residing in storage is met with a greater change in $S_2$ for relatively long sperm in small females (A) but for short sperm in large females (C), with a transitional stage for intermediate female size (B).

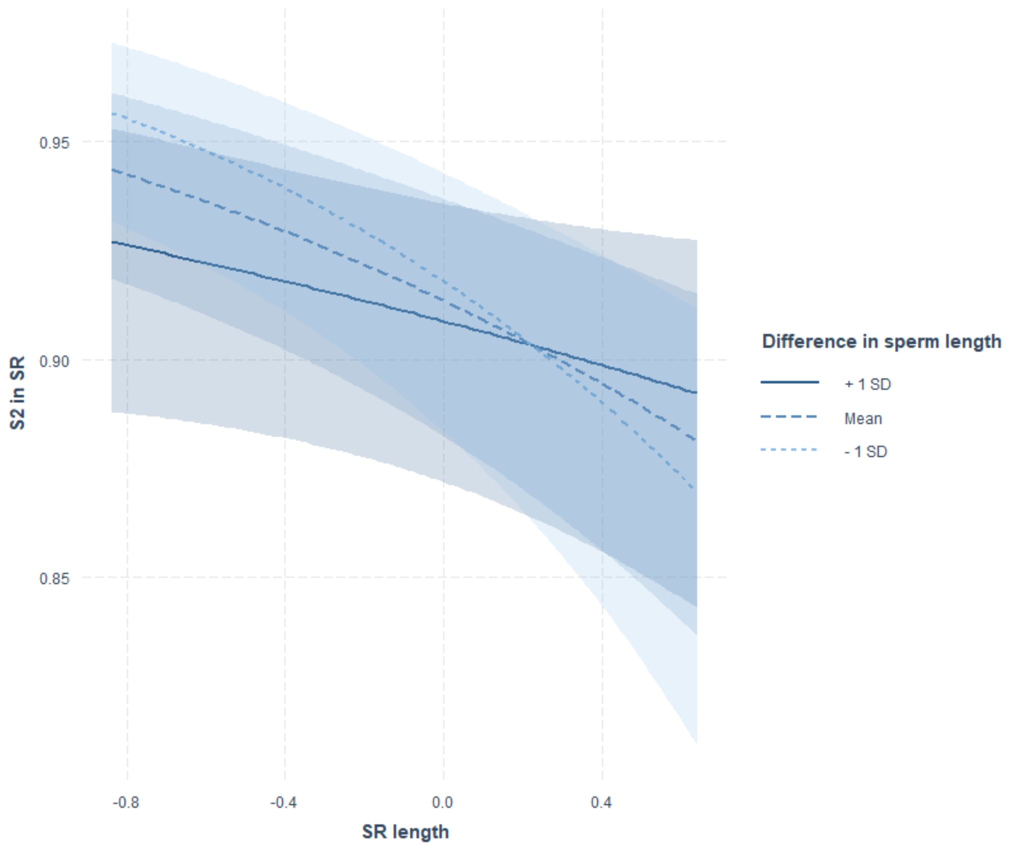


**Supplementary Figure S3:** Conditional effects plot of the two-way interaction between SR length and the difference (2nd male – 1st male) in sperm length explaining variance in the fertilization set ($S_2$ within the female SR). The plot indicates that the second male's sperm representation in the SR declines with any increase in SR length if his sperm are shorter than those of this rival, but less so when he has relatively longer sperm. The shaded areas around lines depict the 95% confidence intervals.

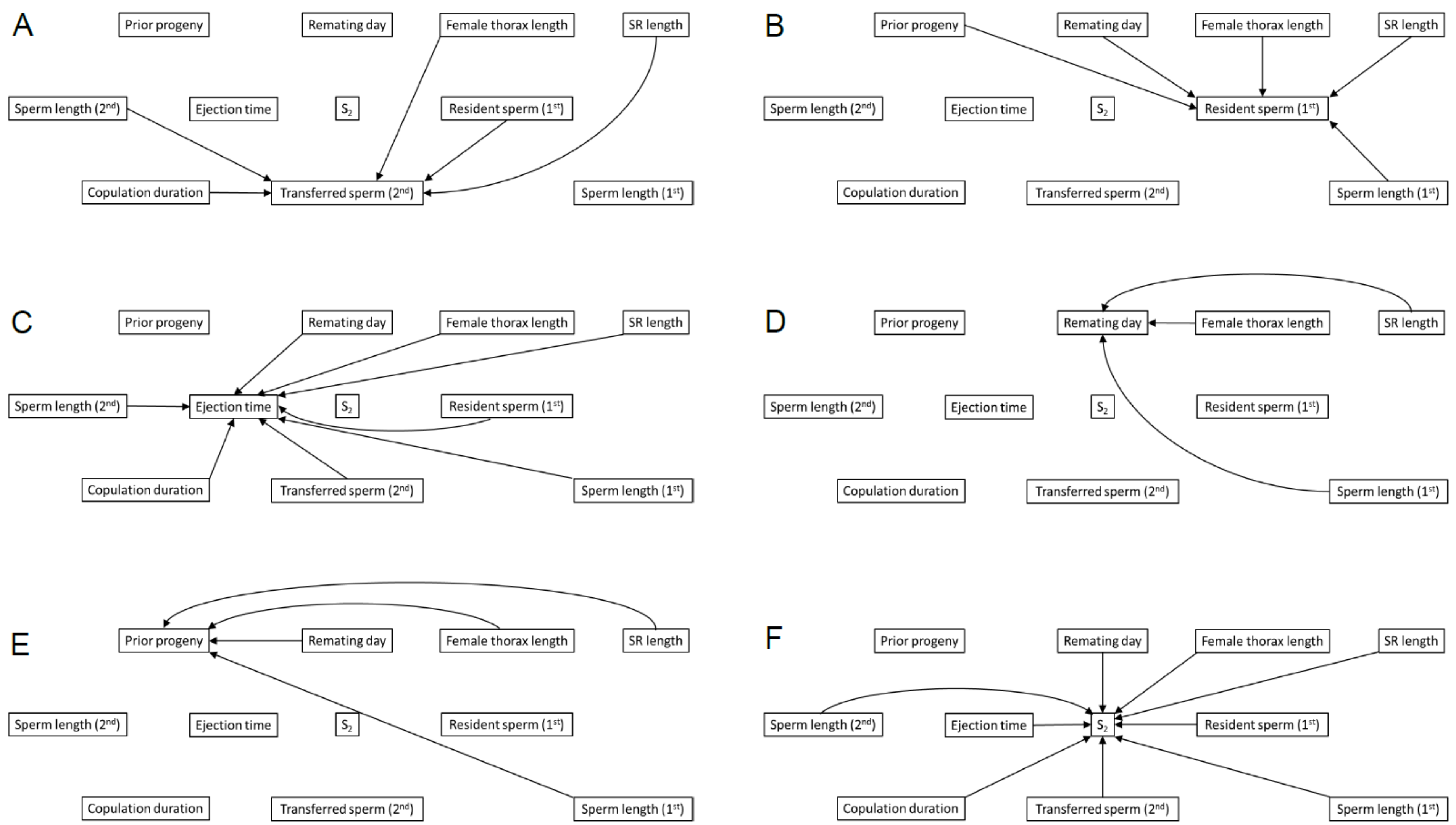


**Supplementary Figure S4:** Visual representation of the GLMM models used in the confirmatory path analysis, with arrows connecting all predictors with the response variable examined. These individual models were then combined to evaluate the entire structural equation model (see Fig. 1 in the main text).

**Supplementary Table S1:** Results of the information-theoretic analyses examining the female ($F_m$), 1st-male ($M_1$) and 2nd-male ($M_2$) genotypic main and interacting effects on the female remating interval ($N$ = 559). Each model was a linear mixed-effects model with the temporal block as a four-level random factor. The last two columns list the results when slightly more complex, nested versions of higher-ranked models are excluded, following the recommendation of (1, 2). The most parsimonious confidence model set is highlighted in bold.

| Int | $F_m$ | $M_1$ | $M_2$ | $F_m \times M_1$ | $F_m \times M_2$ | $M_1 \times M_2$ | $F_m \times M_1 \times M_2$ | *df* | logLik | $AIC_c$ | $\Delta AIC_c$ | $w_i$ | cum $w_i$ | ER | $\Delta AIC_c$ | $w_i$ |
|---|---|---|---|---|---|---|---|---|---|---|---|---|---|---|---|---|
| **2.51** | **+** | | | | | | | **8** | **-714.1** | **1444.5** | **0.00** | **0.614** | **0.614** | **—** | **0.00** | **0.997** |
| 2.47 | + | | + | | | | | 10 | -713.0 | 1446.4 | 1.86 | 0.242 | 0.856 | 2.5 | | |
| 2.69 | + | + | | | | | | 13 | -711.1 | 1448.9 | 4.34 | 0.070 | 0.926 | 8.8 | | |
| 2.49 | + | | + | | + | | | 20 | -704.2 | 1450.0 | 5.50 | 0.039 | 0.965 | 15.7 | | |
| 2.66 | + | + | + | | | | | 15 | -710.0 | 1450.8 | 6.27 | 0.027 | 0.992 | 22.7 | | |
| 2.69 | + | + | + | | + | | | 25 | -701.0 | 1454.4 | 9.87 | 0.004 | 0.996 | 153.5 | | |
| 2.68 | | | | | | | | 3 | -725.0 | 1456.0 | 11.48 | 0.002 | 0.998 | 307.0 | 11.48 | 0.003 |
| 2.66 | | | + | | | | | 5 | -724.1 | 1458.3 | 13.78 | 0.001 | 0.999 | 614.0 | | |
| 2.87 | | + | | | | | | 8 | -722.0 | 1460.2 | 15.64 | 0.000 | 0.999 | ∞ | | |
| 2.85 | | + | + | | | | | 10 | -721.1 | 1462.5 | 18.01 | 0.000 | 0.999 | ∞ | | |
| 2.72 | + | + | + | | | + | | 25 | -708.2 | 1468.8 | 24.28 | 0.000 | 0.999 | ∞ | | |
| 2.75 | + | + | + | | + | + | | 35 | -698.9 | 1472.7 | 28.18 | 0.000 | 0.999 | ∞ | | |
| 2.46 | + | + | | + | | | | 38 | -697.9 | 1477.4 | 32.89 | 0.000 | 0.999 | ∞ | | |
| 2.41 | + | + | + | + | | | | 40 | -696.7 | 1479.6 | 35.12 | 0.000 | 0.999 | ∞ | | |
| 2.92 | | + | + | | | + | | 20 | -719.5 | 1480.5 | 36.01 | 0.000 | 0.999 | ∞ | | |
| 2.45 | + | + | + | + | + | | | 50 | -688.0 | 1486.1 | 41.54 | 0.000 | 0.999 | ∞ | | |
| 2.49 | + | + | + | + | | + | | 50 | -694.7 | 1499.5 | 54.95 | 0.000 | 0.999 | ∞ | | |
| 2.53 | + | + | + | + | + | + | | 60 | -685.8 | 1506.4 | 61.83 | 0.000 | 0.999 | ∞ | | |
| 2.72 | + | + | + | + | + | + | + | 110 | -663.2 | 1600.9 | 156.32 | 0.000 | 0.999 | ∞ | | |
| *SW* | 1.00 | 0.10 | 0.31 | 0.00 | 0.04 | 0.00 | 0.00 | | | | | | | | | |

Int = intercept, (cum) $w_i$ = (cumulative) Akaike weight, ER = evidence ratio (ratio of $w_{best}$ : $w_i$), *SW* = sum of $w_i$ for all models in which the corresponding predictor occurs (note that due to differential representation, *SW* is comparable only among main effects or among two-way interactions but not between main effects and interactions)

**Supplementary Table S2:** Results of the information-theoretic analyses examining the female ($F_m$) and 1st-male ($M_1$) genotypic main and interacting effects on the number of 1st-male sperm residing in the FRT at remating ($N = 567$). Each model was a linear mixed-effects model with the temporal block as a four-level random factor. The most parsimonious confidence model set is highlighted in bold. Note that 2nd males are omitted because they had no effect on the remating interval (Table S1) as the only way they would be able to influence the number of 1st-male sperm remaining in storage.

| Int | $F_m$ | $M_1$ | $F_m \times M_1$ | *df* | logLik | $AIC_c$ | $\Delta AIC_c$ | $w_i$ | cum $w_i$ | ER |
|---|---|---|---|---|---|---|---|---|---|---|
| **278.9** | + | | | **8** | **-3668.1** | **7352.4** | **0.00** | **0.555** | **0.555** | — |
| **270.9** | + | + | | **13** | **-3663.1** | **7352.9** | **0.45** | **0.445** | **1.000** | **1.25** |
| 328.7 | | | | 3 | -3686.9 | 7379.9 | 27.45 | 0.226 | 1.000 | ∞ |
| 318.0 | | + | | 8 | -3682.1 | 7380.4 | 27.99 | 0.205 | 1.000 | ∞ |
| 266.4 | + | + | + | 38 | -3649.6 | 7380.8 | 28.36 | 0.000 | 1.000 | ∞ |
| *SW* | 1.00 | 0.45 | 0.00 | | | | | | | |

Int = intercept, (cum) $w_i$ = (cumulative) Akaike weight, ER = evidence ratio (ratio of $w_{best} : w_i$), *SW* = sum of $w_i$ for all models in which the corresponding predictor occurs (note that due to differential representation, *SW* is comparable only among main effects or among two-way interactions but not between main effects and interactions)

**Supplementary Table S3:** Results of the information-theoretic analyses examining the female ($F_m$) and 1st-male ($M_1$) genotypic main and interacting effects on the number of progeny produced between copulations (*N* = 726). Each model was a linear mixed-effects model with the temporal block as a four-level random factor and controlling for the remating interval (RmI). The last two columns list the results when slightly more complex, nested versions of higher-ranked models are excluded, following the recommendation of (1, 2). The most parsimonious confidence model set is highlighted in bold. Note that 2nd males are omitted because they had no effect on the remating interval (Table S1) as the only way they would be able to influence the number of progeny sired by the first male.

| Int | RmI | $F_m$ | $M_1$ | $F_m \times M_1$ | *df* | logLik | $AIC_c$ | $\Delta AIC_c$ | $w_i$ | cum $w_i$ | ER | $\Delta AIC_c$ | $w_i$ |
|---|---|---|---|---|---|---|---|---|---|---|---|---|---|
| **84.3** | **22.0** | + | + | | **14** | **-3561.3** | **7151.2** | **0.00** | **0.578** | **0.578** | — | **0.00** | **0.578** |
| **80.1** | **22.1** | + | | | **9** | **-3566.8** | **7151.9** | **0.63** | **0.422** | **1.000** | **1.37** | **0.63** | **0.422** |
| 92.4 | 22.4 | + | + | + | 39 | -3548.3 | 7179.1 | 27.86 | 0.000 | 1.000 | ∞ | | |
| 101.2 | 22.1 | | | | 4 | -3616.8 | 7241.7 | 90.42 | 0.000 | 1.000 | ∞ | 90.42 | 0.000 |
| 105.7 | 22.0 | | + | | 9 | -3612.3 | 7242.8 | 91.59 | 0.000 | 1.000 | ∞ | | |
| 83.8 | | + | + | | 13 | -3691.3 | 7409.0 | 257.79 | 0.000 | 1.000 | ∞ | 257.79 | 0.000 |
| 75.9 | | + | | | 8 | -3696.7 | 7409.7 | 258.42 | 0.000 | 1.000 | ∞ | 258.42 | 0.000 |
| 88.1 | | + | + | + | 38 | -3680.4 | 7441.0 | 289.80 | 0.000 | 1.000 | ∞ | | |
| 101.2 | | | | | 3 | -3736.9 | 7479.9 | 328.67 | 0.000 | 1.000 | ∞ | 328.67 | 0.000 |
| 109.2 | | | + | | 8 | -3732.0 | 7480.3 | 329.04 | 0.000 | 1.000 | ∞ | | |
| *SW* | 1.00 | 1.00 | 0.58 | 0.00 | | | | | | | | | |

Int = intercept, (cum) $w_i$ = (cumulative) Akaike weight, ER = evidence ratio (ratio of $w_{best} : w_i$), *SW* = sum of $w_i$ for all models in which the corresponding predictor occurs (note that due to differential representation, *SW* is comparable only among main effects or among two-way interactions but not between main effects and interactions)

**Supplementary Table S4:** Results of the information-theoretic analyses examining the female ($F_m$), 1st-male ($M_1$) and 2nd-male ($M_2$) genotypic main and interacting effects on the duration of the second mating ($N$ = 761). Each model was a linear mixed-effects model with the temporal block as a four-level random factor. The last two columns list the results when slightly more complex, nested versions of higher-ranked models are excluded, following the recommendation of (1, 2). The most parsimonious confidence model set is highlighted in bold.

| Int | $F_m$ | $M_1$ | $M_2$ | $F_m \times M_1$ | $F_m \times M_2$ | $M_1 \times M_2$ | $F_m \times M_1 \times M_2$ | *df* | logLik | $AIC_c$ | $\Delta AIC_c$ | $w_i$ | cum $w_i$ | ER | $\Delta AIC_c$ | $w_i$ |
|---|---|---|---|---|---|---|---|---|---|---|---|---|---|---|---|---|
| **27.5** | | | + | | | | | **5** | **-2592.8** | **5195.6** | **0.00** | **0.897** | **0.897** | **—** | **0.00** | **1.000** |
| 27.9 | + | | + | | | | | 10 | -2590.1 | 5200.4 | 4.83 | 0.080 | 0.977 | 11.21 | | |
| 27.7 | | + | + | | | | | 10 | -2591.4 | 5203.1 | 7.50 | 0.021 | 0.998 | 42.71 | | |
| 28.1 | + | + | + | | | | | 15 | -2588.6 | 5207.9 | 12.35 | 0.002 | 1.000 | 448.50 | | |
| 28.3 | + | | + | | + | | | 20 | -2585.0 | 5211.2 | 15.65 | 0.000 | 1.000 | ∞ | | |
| 27.7 | | + | + | | | + | | 20 | -2588.2 | 5217.4 | 21.86 | 0.000 | 1.000 | ∞ | | |
| 28.5 | + | + | + | | + | | | 25 | -2583.6 | 5219.0 | 23.44 | 0.000 | 1.000 | ∞ | | |
| 28.4 | + | + | + | + | | | | 40 | -2568.8 | 5222.2 | 26.63 | 0.000 | 1.000 | ∞ | | |
| 28.1 | + | + | + | | | + | | 25 | -2585.4 | 5222.5 | 26.94 | 0.000 | 1.000 | ∞ | | |
| 29.0 | + | + | + | + | + | | | 50 | -2563.0 | 5233.2 | 37.62 | 0.000 | 1.000 | ∞ | | |
| 28.5 | + | + | + | | + | + | | 35 | -2580.4 | 5234.3 | 38.68 | 0.000 | 1.000 | ∞ | | |
| 28.5 | + | + | + | + | | + | | 50 | -2565.3 | 5237.7 | 42.10 | 0.000 | 1.000 | ∞ | | |
| 29.1 | + | + | + | + | + | + | | 60 | -2559.4 | 5249.3 | 53.73 | 0.000 | 1.000 | ∞ | | |
| 28.4 | | | | | | | | 3 | -2645.1 | 5296.3 | 100.69 | 0.000 | 1.000 | ∞ | 100.69 | 0.000 |
| 28.8 | + | | | | | | | 8 | -2642.7 | 5301.6 | 105.99 | 0.000 | 1.000 | ∞ | | |
| 28.6 | | + | | | | | | 8 | -2643.9 | 5304.1 | 108.47 | 0.000 | 1.000 | ∞ | | |
| 29.0 | + | + | | | | | | 13 | -2641.4 | 5309.4 | 113.79 | 0.000 | 1.000 | ∞ | | |
| 27.5 | + | + | + | + | + | + | + | 110 | -2528.3 | 5314.1 | 118.55 | 0.000 | 1.000 | ∞ | | |
| 29.2 | + | + | | + | | | | 38 | -2623.6 | 5327.2 | 131.66 | 0.000 | 1.000 | ∞ | | |
| *SW* | 0.08 | 0.02 | 1.00 | 0.00 | 0.00 | 0.00 | 0.00 | | | | | | | | | |

Int = intercept, (cum) $w_i$ = (cumulative) Akaike weight, ER = evidence ratio (ratio of $w_{best}$ : $w_i$), *SW* = sum of $w_i$ for all models in which the corresponding predictor occurs (note that due to differential representation, *SW* is comparable only among main effects or among two-way interactions but not between main effects and interactions)

**Supplementary Table S5:** Results of the information-theoretic analyses examining the female ($F_m$), 1st-male ($M_1$) and 2nd-male ($M_2$) genotypic main and interacting effects on the number of sperm transferred by the second male ($N = 559$). Each model was a linear mixed-effects model with the temporal block as a four-level random factor. The last two columns list the results when slightly more complex, nested versions of higher-ranked models are excluded, following the recommendation of (1, 2). The most parsimonious confidence model set is highlighted in bold.

| **Int** | **$F_m$** | **$M_1$** | **$M_2$** | **$F_m \times M_1$** | **$F_m \times M_2$** | **$M_1 \times M_2$** | **$F_m \times M_1 \times M_2$** | ***df*** | **logLik** | **$AIC_c$** | **$\Delta AIC_c$** | **$w_i$** | **cum $w_i$** | **ER** | **$\Delta AIC_c$** | **$w_i$** |
|---|---|---|---|---|---|---|---|---|---|---|---|---|---|---|---|---|
| **1387** | | | **+** | | | | | **5** | **-4150.7** | **8311.5** | **0.00** | **0.847** | **0.847** | **—** | **0.00** | **0.998** |
| 1380 | + | | + | | | | | 10 | -4147.6 | 8315.6 | 4.10 | 0.109 | 0.956 | 7.77 | | |
| 1409 | | + | + | | | | | 10 | -4148.8 | 8318.0 | 6.51 | 0.033 | 0.989 | 25.67 | | |
| 1543 | | + | + | | | + | | 20 | -4140.2 | 8322.0 | 10.49 | 0.004 | 0.993 | 211.75 | | |
| 1403 | + | + | + | | | | | 15 | -4145.8 | 8322.4 | 10.97 | 0.004 | 0.997 | 211.75 | | |
| 1292 | | | | | | | | 3 | -4158.8 | 8323.6 | 12.16 | 0.002 | 0.999 | 423.50 | 12.16 | 0.003 |
| 1541 | + | + | + | | | + | | 25 | -4137.1 | 8326.6 | 15.12 | 0.000 | 0.999 | ∞ | | |
| 1282 | + | | | | | | | 8 | -4155.6 | 8327.5 | 16.01 | 0.000 | 0.999 | ∞ | | |
| 1320 | | + | | | | | | 8 | -4156.6 | 8329.4 | 17.95 | 0.000 | 0.999 | ∞ | | |
| 1352 | + | | + | | + | | | 20 | -4144.1 | 8329.9 | 18.39 | 0.000 | 0.999 | ∞ | | |
| 1311 | + | + | | | | | | 13 | -4153.5 | 8333.7 | 22.20 | 0.000 | 0.999 | ∞ | | |
| 1382 | + | + | + | | + | | | 25 | -4142.3 | 8337.0 | 25.50 | 0.000 | 0.999 | ∞ | | |
| 1524 | + | + | + | | + | + | | 35 | -4133.9 | 8342.6 | 31.18 | 0.000 | 0.999 | ∞ | | |
| 1432 | + | + | + | + | | | | 40 | -4129.4 | 8345.1 | 33.66 | 0.000 | 0.999 | ∞ | | |
| 1575 | + | + | + | + | | + | | 50 | -4120.9 | 8351.8 | 40.30 | 0.000 | 0.999 | ∞ | | |
| 1332 | + | + | | + | | | | 38 | -4137.2 | 8356.2 | 44.71 | 0.000 | 0.999 | ∞ | | |
| 1411 | + | + | + | + | + | | | 50 | -4125.9 | 8361.8 | 50.31 | 0.000 | 0.999 | ∞ | | |
| 1558 | + | + | + | + | + | + | | 60 | -4117.7 | 8370.1 | 58.68 | 0.000 | 0.999 | ∞ | | |
| 1723 | + | + | + | + | + | + | + | 110 | -4077.0 | 8428.6 | 117.08 | 0.000 | 0.999 | ∞ | | |
| *SW* | 0.11 | 0.04 | 1.00 | 0.00 | 0.00 | 0.00 | 0.00 | | | | | | | | | |

Int = intercept, (cum) $w_i$ = (cumulative) Akaike weight, ER = evidence ratio (ratio of $w_{best} : w_i$), *SW* = sum of $w_i$ for all models in which the corresponding predictor occurs (note that due to differential representation, *SW* is comparable only among main effects or among two-way interactions but not between main effects and interactions)

**Supplementary Table S6:** Results of the information-theoretic analyses examining the female ($F_m$), 1st-male ($M_1$) and 2nd-male ($M_2$) genotypic main and interacting effects on the time to female sperm ejection (*N* = 664). Each model was a linear mixed-effects model with the temporal block as a four-level random factor. Ejection time was log-transformed to normalize the data distribution. The last two columns list the results when slightly more complex, nested versions of higher-ranked models are excluded, following the recommendation of (1, 2). The most parsimonious confidence model set is highlighted in bold.

| Int | $F_m$ | $M_1$ | $M_2$ | $F_m \times M_1$ | $F_m \times M_2$ | $M_1 \times M_2$ | $F_m \times M_1 \times M_2$ | *df* | logLik | $AIC_c$ | $\Delta AIC_c$ | $w_i$ | cum $w_i$ | ER | $\Delta AIC_c$ | $w_i$ |
|---|---|---|---|---|---|---|---|---|---|---|---|---|---|---|---|---|
| **4.56** | + | | + | | + | | | **20** | **-337.9** | **717.1** | **0.00** | **0.888** | **0.888** | — | **0.00** | **0.898** |
| **4.53** | + | | + | | | | | **10** | **-350.6** | **721.6** | **4.56** | **0.091** | **0.979** | **9.76** | **4.56** | **0.092** |
| 4.53 | + | | | | | | | 8 | -354.9 | 726.0 | 8.92 | 0.010 | 0.989 | 88.80 | 8.92 | 0.010 |
| 4.54 | + | + | + | | + | | | 25 | -337.0 | 726.1 | 9.05 | 0.010 | 0.999 | 88.80 | | |
| 4.52 | + | + | + | | | | | 15 | -349.7 | 730.0 | 12.98 | 0.001 | 1.000 | 888.00 | | |
| 4.51 | + | + | | | | | | 13 | -353.9 | 734.3 | 17.22 | 0.000 | 1.000 | ∞ | | |
| 4.46 | + | + | + | | + | + | | 35 | -332.4 | 738.7 | 21.65 | 0.000 | 1.000 | ∞ | | |
| 4.43 | + | + | + | | | + | | 25 | -345.5 | 743.1 | 26.02 | 0.000 | 1.000 | ∞ | | |
| 4.62 | + | + | + | + | + | | | 50 | -324.2 | 756.8 | 39.71 | 0.000 | 1.000 | ∞ | | |
| 4.58 | + | + | + | + | | | | 40 | -336.9 | 759.1 | 42.07 | 0.000 | 1.000 | ∞ | | |
| 4.57 | + | + | | + | | | | 38 | -341.0 | 762.7 | 45.63 | 0.000 | 1.000 | ∞ | | |
| 4.53 | + | + | + | + | + | + | | 60 | -319.1 | 770.3 | 53.25 | 0.000 | 1.000 | ∞ | | |
| 4.49 | + | + | + | + | | + | | 50 | -332.4 | 773.2 | 56.09 | 0.000 | 1.000 | ∞ | | |
| 4.37 | | | + | | | | | 5 | -400.1 | 810.3 | 93.24 | 0.000 | 1.000 | ∞ | 93.24 | 0.000 |
| 4.36 | | | | | | | | 3 | -403.1 | 812.3 | 95.21 | 0.000 | 1.000 | ∞ | 95.21 | 0.000 |
| 4.34 | | + | + | | | | | 10 | -398.9 | 818.2 | 101.13 | 0.000 | 1.000 | ∞ | | |
| 4.33 | | + | | | | | | 8 | -401.9 | 820.1 | 103.00 | 0.000 | 1.000 | ∞ | | |
| 4.25 | | + | + | | | + | | 20 | -395.3 | 831.9 | 114.84 | 0.000 | 1.000 | ∞ | | |
| 4.52 | + | + | + | + | + | + | + | 110 | -297.8 | 859.7 | 142.64 | 0.000 | 1.000 | ∞ | | |
| *SW* | 1.00 | 0.01 | 0.99 | 0.00 | 0.90 | 0.00 | 0.00 | | | | | | | | | |

Int = intercept, (cum) $w_i$ = (cumulative) Akaike weight, ER = evidence ratio (ratio of $w_{best}$ : $w_i$), *SW* = sum of $w_i$ for all models in which the corresponding predictor occurs (note that due to differential representation, *SW* is comparable only among main effects or among two-way interactions but not between main effects and interactions)

**Supplementary Table S7:** Results of the information-theoretic analyses examining the female ($F_m$), 1st-male ($M_1$) and 2nd-male ($M_2$) genotypic main and interacting effects on total $S_2$ after female sperm ejection ($N = 583$). Each model was a generalized linear mixed-effects model with the temporal block as a four-level random factor and an observation-level random effect to account of overdispersion (final dispersion = 1.08). The last two columns list the results when slightly more complex, nested versions of higher-ranked models are excluded, following the recommendation of (1, 2). The most parsimonious confidence model set is highlighted in bold.

| Int | $F_m$ | $M_1$ | $M_2$ | $F_m \times M_1$ | $F_m \times M_2$ | $M_1 \times M_2$ | $F_m \times M_1 \times M_2$ | *df* | logLik | $AIC_c$ | $\Delta AIC_c$ | $w_i$ | cum $w_i$ | ER | $\Delta AIC_c$ | $w_i$ |
|---|---|---|---|---|---|---|---|---|---|---|---|---|---|---|---|---|
| **1.47** | **+** | **+** | **+** | | | | | **15** | **-3436.4** | **6903.6** | **0.00** | **0.996** | **0.996** | **—** | **0.00** | **0.999** |
| 1.55 | + | + | + | | + | | | 25 | -3431.8 | 6915.9 | 12.27 | 0.002 | 0.998 | 498.00 | | |
| 1.63 | + | | + | | | | | 10 | -3448.3 | 6917.0 | 13.42 | 0.001 | 0.999 | 996.00 | 13.42 | 0.001 |
| 1.41 | + | + | + | | | + | | 25 | -3433.1 | 6918.5 | 14.88 | 0.001 | 1.000 | 996.00 | | |
| 1.43 | | + | + | | | | | 10 | -3450.6 | 6921.6 | 18.05 | 0.000 | 1.000 | ∞ | 18.05 | 0.000 |
| 1.19 | + | + | | | | | | 13 | -3447.8 | 6922.2 | 18.62 | 0.000 | 1.000 | ∞ | 18.62 | 0.000 |
| 1.83 | + | + | + | + | | | | 40 | -3418.9 | 6923.8 | 20.19 | 0.000 | 1.000 | ∞ | | |
| 1.70 | + | | + | | + | | | 20 | -3443.7 | 6928.9 | 25.34 | 0.000 | 1.000 | ∞ | | |
| 1.51 | + | + | + | | + | + | | 35 | -3428.5 | 6931.6 | 28.03 | 0.000 | 1.000 | ∞ | | |
| 1.34 | + | | | | | | | 8 | -3458.9 | 6934.0 | 30.40 | 0.000 | 1.000 | ∞ | 30.40 | 0.000 |
| 1.61 | | | + | | | | | 5 | -3462.0 | 6934.1 | 30.52 | 0.000 | 1.000 | ∞ | 30.52 | 0.000 |
| 1.37 | | + | + | | | + | | 20 | -3447.1 | 6935.6 | 32.03 | 0.000 | 1.000 | ∞ | | |
| 1.91 | + | + | + | + | + | | | 50 | -3413.7 | 6936.9 | 33.31 | 0.000 | 1.000 | ∞ | | |
| 1.77 | + | + | + | + | | + | | 50 | -3415.0 | 6939.5 | 35.92 | 0.000 | 1.000 | ∞ | | |
| 1.16 | | + | | | | | | 8 | -3461.7 | 6939.6 | 35.97 | 0.000 | 1.000 | ∞ | 35.97 | 0.000 |
| 1.54 | + | + | | + | | | | 38 | -3430.7 | 6942.8 | 39.25 | 0.000 | 1.000 | ∞ | | |
| 1.32 | | | | | | | | 3 | -3472.3 | 6950.6 | 47.00 | 0.000 | 1.000 | ∞ | 47.00 | 0.000 |
| 1.87 | + | + | + | + | + | + | | 60 | -3409.7 | 6953.5 | 49.93 | 0.000 | 1.000 | ∞ | | |
| 1.67 | + | + | + | + | + | + | + | 110 | -3380.8 | 7033.3 | 129.76 | 0.000 | 1.000 | ∞ | | |
| *SW* | 1.00 | 1.00 | 1.00 | 0.00 | 0.00 | 0.00 | 0.00 | | | | | | | | | |

Int = intercept, (cum) $w_i$ = (cumulative) Akaike weight, ER = evidence ratio (ratio of $w_{best} : w_i$), *SW* = sum of $w_i$ for all models in which the corresponding predictor occurs (note that due to differential representation, *SW* is comparable only among main effects or among two-way interactions but not between main effects and interactions)

**Supplementary Table S8:** Results of the information-theoretic analyses examining the female ($F_m$), 1st-male ($M_1$) and 2nd-male ($M_2$) genotypic main and interacting effects on $S_2$ within the seminal receptacle (i.e., the “fertilization set”) after female sperm ejection ($N$ = 598). Each model was a generalized linear mixed-effects model with the temporal block as a four-level random factor and an observation-level random effect to account of overdispersion (final dispersion = 1.08). The last two columns list the results when slightly more complex, nested versions of higher-ranked models are excluded, following the recommendation of (1, 2). The most parsimonious confidence model set is highlighted in bold.

| Int | $F_m$ | $M_1$ | $M_2$ | $F_m \times M_1$ | $F_m \times M_2$ | $M_1 \times M_2$ | $F_m \times M_1 \times M_2$ | *df* | logLik | $AIC_c$ | $\Delta AIC_c$ | $w_i$ | cum $w_i$ | ER | $\Delta AIC_c$ | $w_i$ |
|---|---|---|---|---|---|---|---|---|---|---|---|---|---|---|---|---|
| **3.16** | **+** | | **+** | | | | | **10** | **-2941.0** | **5902.4** | **0.00** | **0.937** | **0.937** | **—** | **0.00** | **0.999** |
| 3.32 | + | + | + | | | | | 15 | -2938.7 | 5908.3 | 5.87 | 0.050 | 0.987 | 18.83 | | |
| 3.09 | + | | + | | + | | | 20 | -2934.9 | 5911.3 | 8.87 | 0.011 | 0.998 | 84.37 | | |
| 2.63 | | | + | | | | | 5 | -2953.4 | 5916.9 | 14.49 | 0.001 | 0.999 | 1399.06 | 14.49 | 0.001 |
| 3.24 | + | + | + | | + | | | 25 | -2932.6 | 5917.5 | 15.09 | 0.000 | 0.999 | ∞ | | |
| 4.56 | + | + | + | + | | | | 40 | -2916.4 | 5918.7 | 16.28 | 0.000 | 1.000 | ∞ | | |
| 2.82 | + | | | | | | | 8 | -2951.6 | 5919.4 | 16.93 | 0.000 | 1.000 | ∞ | 16.93 | 0.000 |
| 2.97 | + | + | + | | | + | | 25 | -2934.7 | 5921.6 | 19.19 | 0.000 | 1.000 | ∞ | | |
| 2.80 | | + | + | | | | | 10 | -2950.8 | 5922.1 | 19.64 | 0.000 | 1.000 | ∞ | | |
| 3.02 | + | + | | | | | | 13 | -2949.2 | 5924.9 | 22.52 | 0.000 | 1.000 | ∞ | | |
| 4.47 | + | + | + | + | + | | | 50 | -2910.1 | 5929.6 | 27.18 | 0.000 | 1.000 | ∞ | | |
| 2.89 | + | + | + | | + | + | | 35 | -2928.7 | 5931.8 | 29.38 | 0.000 | 1.000 | ∞ | | |
| 2.30 | | | | | | | | 3 | -2963.7 | 5933.3 | 30.93 | 0.000 | 1.000 | ∞ | 30.93 | 0.000 |
| 4.23 | + | + | + | + | | + | | 50 | -2912.4 | 5934.2 | 31.77 | 0.000 | 1.000 | ∞ | | |
| 2.44 | | + | + | | | + | | 20 | -2947.0 | 5935.5 | 33.07 | 0.000 | 1.000 | ∞ | | |
| 4.28 | + | + | | + | | | | 38 | -2927.1 | 5935.6 | 33.17 | 0.000 | 1.000 | ∞ | | |
| 2.49 | | + | | | | | | 8 | -2961.0 | 5938.3 | 35.87 | 0.000 | 1.000 | ∞ | | |
| 4.15 | + | + | + | + | + | + | | 60 | -2906.3 | 5946.3 | 43.84 | 0.000 | 1.000 | ∞ | | |
| 3.77 | + | + | + | + | + | + | + | 110 | -2879.0 | 6028.2 | 125.77 | 0.000 | 1.000 | ∞ | | |
| *SW* | 1.00 | 0.05 | 1.00 | 0.00 | 0.01 | 0.00 | 0.00 | | | | | | | | | |

Int = intercept, (cum) $w_i$ = (cumulative) Akaike weight, ER = evidence ratio (ratio of $w_{best}$ : $w_i$), *SW* = sum of $w_i$ for all models in which the corresponding occurs (note that due to differential representation, *SW* is comparable only among main effects or among two-way interactions but not between main effects and interactions)

**Supplementary Table S9:** Results of the information-theoretic analyses examining the effects of copulation duration (C), the number of 1st-male sperm residing in the FRT at remating (R) and female thorax length (T) on the number of sperm transferred by the second male (*N* = 558). Each model was a linear mixed-effects model with the temporal block as a four-level random factor. The last two columns list the results when slightly more complex, nested versions of higher-ranked models are excluded, following the recommendation of (1, 2). The most parsimonious confidence model set is highlighted in bold.

| Model | *df* | logLik | AIC$_c$ | ΔAIC$_c$ | $w_i$ | cum $w_i$ | ER | ΔAIC$_c$ | $w_i$ |
|---|---|---|---|---|---|---|---|---|---|
| **C + R + C×R** | **10** | **-734.19** | **1488.78** | **0.00** | **0.30** | **0.30** | **—** | **0.00** | **0.30** |
| **C + R** | **9** | **-735.55** | **1489.42** | **0.64** | **0.23** | **0.53** | **1.30** | **0.64** | **0.23** |
| C + R + T + C×R | 11 | -734.01 | 1490.51 | 1.72 | 0.13 | 0.66 | 2.31 | | |
| C + R + T | 10 | -735.36 | 1491.13 | 2.34 | 0.09 | 0.75 | 3.33 | | |
| C + R + T + C×R + R×T | 12 | -733.71 | 1492.00 | 3.21 | 0.06 | 0.81 | 5.00 | | |
| C + R + T + C×R + C×T | 12 | -733.84 | 1492.26 | 3.48 | 0.05 | 0.86 | 6.00 | | |
| C + R + T + R×T | 11 | -735.06 | 1492.60 | 3.82 | 0.04 | 0.90 | 7.50 | | |
| C + R + T + C×T | 11 | -735.14 | 1492.77 | 3.99 | 0.04 | 0.94 | 7.50 | | |
| C + R + T + C×R + C×T + R×T | 13 | -733.55 | 1493.76 | 4.98 | 0.02 | 0.96 | 15.00 | | |
| C + R + T + C×T + R×T | 12 | -734.84 | 1494.26 | 5.47 | 0.03 | 0.99 | 10.00 | | |
| C + R + T + C×R + C×T + R×T + C×R×T | 14 | -733.29 | 1495.35 | 6.57 | 0.01 | 1.00 | 30.00 | | |
| R | 8 | -740.51 | 1497.29 | 8.50 | 0.00 | 1.00 | ∞ | 8.50 | 0.00 |
| R + T | 9 | -740.39 | 1499.11 | 10.32 | 0.00 | 1.00 | ∞ | | |
| R + T + R×T | 10 | -740.09 | 1500.58 | 11.79 | 0.00 | 1.00 | ∞ | | |
| C | 8 | -753.37 | 1523.01 | 34.22 | 0.00 | 1.00 | ∞ | 34.22 | 0.00 |
| C + T | 9 | -753.36 | 1525.05 | 36.27 | 0.00 | 1.00 | ∞ | | |
| C + T + C×T | 10 | -753.14 | 1526.67 | 37.89 | 0.00 | 1.00 | ∞ | | |
| (Null) | 7 | -757.47 | 1529.14 | 40.35 | 0.00 | 1.00 | ∞ | 40.35 | 0.00 |
| T | 8 | -757.47 | 1531.19 | 42.41 | 0.00 | 1.00 | ∞ | | |

Coefficients of the above analyses (naturally) averaged across the reduced confidence model set, including the conditional standard errors (SE) and 95% confidence limits (CL), the relative variable importance (*RI*) and number of models including each term (*N*).

| | Full confidence set | | | | | Excluding nested models | | | | |
|---|---|---|---|---|---|---|---|---|---|---|
| **Parameter** | **Estimate** | **SE** | **95% CL** | ***RI*** | ***N*** | **Estimate** | **SE** | **95% CL** | ***RI*** | ***N*** |
| (Intercept) | 0.01 | 0.20 | (-0.37, 0.40) | | 10 | 0.01 | 0.20 | (-0.37, 0.40) | | 2 |
| **R** | **0.48** | **0.08** | **(0.32, 0.63)** | **1.00** | **10** | **0.48** | **0.08** | **(0.32, 0.63)** | **1.00** | **2** |
| **C** | **0.26** | **0.08** | **(0.10, 0.42)** | **1.00** | **10** | **0.26** | **0.08** | **(0.10, 0.42)** | **1.00** | **2** |
| C×R | 0.25 | 0.15 | (-0.05, 0.55) | 0.58 | 5 | 0.25 | 0.15 | (-0.05, 0.55) | 0.58 | 1 |
| T | 0.07 | 0.11 | (-0.15, 0.28) | 0.47 | 8 | | | | | |
| R×T | -0.12 | 0.16 | (-0.43, 0.19) | 0.15 | 4 | | | | | |
| C×T | -0.10 | 0.16 | (-0.42, 0.22) | 0.14 | 4 | | | | | |

**Supplementary Table S10:** Results of the information-theoretic analyses examining the effects of the difference in sperm length (L), the difference in sperm number (N) and seminal receptacle length (S) on the time to female sperm ejection (*N* = 529). Each model was a linear mixed-effects model with the temporal block as a four-level random factor. The last two columns list the results when slightly more complex, nested versions of higher-ranked models are excluded, following the recommendation of (1, 2). The most parsimonious confidence model set is highlighted in bold.

| Model | *df* | logLik | $AIC_c$ | $\Delta AIC_c$ | $w_i$ | cum $w_i$ | ER | $\Delta AIC_c$ | $w_i$ |
|---|---|---|---|---|---|---|---|---|---|
| **L** | **8** | **-711.91** | **1440.10** | **0.00** | **0.23** | **0.23** | **—** | **0.00** | **0.61** |
| **(Null)** | **7** | **-713.39** | **1441.00** | **0.90** | **0.15** | **0.38** | **1.53** | **0.90** | **0.39** |
| L + S + L×S | 10 | -710.62 | 1441.67 | 1.57 | 0.11 | 0.49 | 2.09 | | |
| L + S | 9 | -711.83 | 1442.01 | 1.91 | 0.09 | 0.58 | 2.56 | | |
| L + N | 9 | -711.88 | 1442.11 | 2.01 | 0.08 | 0.66 | 2.88 | | |
| S | 8 | -713.31 | 1442.90 | 2.80 | 0.06 | 0.72 | 3.83 | | |
| N | 8 | -713.36 | 1443.00 | 2.90 | 0.05 | 0.77 | 4.60 | | |
| L + N + L×N | 10 | -711.46 | 1443.35 | 3.25 | 0.05 | 0.82 | 4.60 | | |
| L + N + S + L×S | 11 | -710.57 | 1443.65 | 3.55 | 0.04 | 0.86 | 5.75 | | |
| L + N + S | 10 | -711.80 | 1444.02 | 3.92 | 0.03 | 0.89 | 7.67 | | |
| N + S | 9 | -713.28 | 1444.90 | 4.80 | 0.02 | 0.91 | 11.50 | | |
| L + N + S + L×N + L×S | 12 | -710.21 | 1445.02 | 4.92 | 0.02 | 0.93 | 11.50 | | |
| L + N + S + L×N | 11 | -711.38 | 1445.28 | 5.18 | 0.02 | 0.95 | 11.50 | | |
| L + N + S + L×S + N×S | 12 | -710.51 | 1445.63 | 5.53 | 0.01 | 0.96 | 23.00 | | |
| L + N + S + N×S | 11 | -711.74 | 1445.98 | 5.88 | 0.01 | 0.97 | 23.00 | | |
| N + S + N×S | 10 | -713.22 | 1446.87 | 6.77 | 0.01 | 0.98 | 23.00 | | |
| L + N + S + L×N + L×S + N×S | 13 | -710.13 | 1446.96 | 6.86 | 0.01 | 0.99 | 23.00 | | |
| L + N + S + L×N +N×S | 12 | -711.30 | 1447.20 | 7.10 | 0.01 | 1.00 | 23.00 | | |
| L + N + S + L×N + L×S + N×S + L×N×S | 14 | -709.68 | 1448.18 | 8.08 | 0.00 | 1.00 | ∞ | | |

Coefficients of the above analyses (naturally) averaged across the reduced confidence model set, including the conditional standard errors (SE) and 95% confidence limits (CL), the relative variable importance (*RI*) and number of models including each term (*N*).

| | Full confidence set | | | | | Excluding nested models | | | | |
|---|---|---|---|---|---|---|---|---|---|---|
| **Parameter** | **Estimate** | **SE** | **95% CL** | ***RI*** | ***N*** | **Estimate** | **SE** | **95% CL** | ***RI*** | ***N*** |
| (Intercept) | -0.02 | 0.18 | (-0.37, 0.34) | NA | 15 | -0.02 | 0.19 | (-0.38, 0.35) | 0.08 | 0.94 |
| **L** | **0.20** | **0.08** | **(0.05, 0.36)** | **0.71** | **11** | **0.20** | **0.08** | **(0.05, 0.36)** | **2.56** | **0.01** |
| S | -0.13 | 0.32 | (-0.75, 0.49) | 0.42 | 10 | | | | | |
| N | -0.03 | 0.09 | (-0.19, 0.14) | 0.35 | 4 | | | | | |
| L×S | 0.25 | 0.16 | (-0.07, 0.56) | 0.18 | 10 | | | | | |
| L×N | -0.15 | 0.17 | (-0.48, 0.18) | 0.09 | 3 | | | | | |
| N×S | -0.05 | 0.16 | (-0.36, 0.25) | 0.03 | 2 | | | | | |

**Supplementary Table S11:** Results of the information-theoretic analyses examining the effects of the difference in sperm length (L), the difference in sperm number (N) and the time to female sperm ejection (E) on the relative numbers of sperm stored between males (i.e., total $S_2$, $N = 505$). Each model was a GLMM with the temporal block as a four-level random factor. An observation-level random effect was included to account for overdispersion. The last two columns list the results when slightly more complex, nested versions of higher-ranked models are excluded, following the recommendation of (1, 2). The most parsimonious confidence model set is highlighted in bold.

| Model | *df* | logLik | $AIC_c$ | $\Delta AIC_c$ | $w_i$ | cum $w_i$ | ER | $\Delta AIC_c$ | $w_i$ |
|---|---|---|---|---|---|---|---|---|---|
| **N + E + N×E** | **10** | **-2952.99** | **5926.42** | **0.00** | **0.38** | 0.38 | — | **0.00** | **0.95** |
| L + N + E + L×N + N×E | 12 | -2951.51 | 5927.65 | 1.22 | 0.21 | 0.59 | 1.81 | | |
| L + N + E + N×E | 11 | -2952.60 | 5927.74 | 1.32 | 0.20 | 0.79 | 1.90 | | |
| L + N + E + L×N + L×E + N×E | 13 | -2951.48 | 5929.70 | 3.28 | 0.07 | 0.86 | 5.43 | | |
| L + N + E + L×E + N×E + | 12 | -2952.58 | 5929.79 | 3.37 | 0.07 | 0.93 | 5.43 | | |
| L + N + E + L×N + L×E + N×E + L×N×E | 14 | -2951.22 | 5931.30 | 4.88 | 0.04 | 0.97 | 9.50 | | |
| L + N + E + L×N | 11 | -2955.44 | 5933.41 | 6.99 | 0.02 | 0.99 | 19.00 | 6.99 | 0.03 |
| N + E | 9 | -2957.68 | 5933.73 | 7.30 | 0.01 | 1.00 | 38.00 | 7.30 | 0.02 |
| L + N + E | 10 | -2957.38 | 5935.20 | 8.78 | 0.00 | 1.00 | ∞ | | |
| L + N + E + L×N + L×E | 12 | -2955.42 | 5935.47 | 9.04 | 0.00 | 1.00 | ∞ | | |
| L + N + E + L×E | 11 | -2957.36 | 5937.26 | 10.83 | 0.00 | 1.00 | ∞ | | |
| N | 8 | -2971.05 | 5958.38 | 31.96 | 0.00 | 1.00 | ∞ | 31.96 | 0.00 |
| L + N + L×N | 10 | -2968.98 | 5958.41 | 31.99 | 0.00 | 1.00 | ∞ | | |
| L + N | 9 | -2970.56 | 5959.48 | 33.05 | 0.00 | 1.00 | ∞ | | |
| E | 8 | -2986.81 | 5989.90 | 63.48 | 0.00 | 1.00 | ∞ | 63.48 | 0.00 |
| L + E | 9 | -2986.75 | 5991.87 | 65.45 | 0.00 | 1.00 | ∞ | | |
| L + E + L×E | 10 | -2986.75 | 5993.95 | 67.53 | 0.00 | 1.00 | ∞ | | |
| (Null) | 7 | -2998.32 | 6010.87 | 84.45 | 0.00 | 1.00 | ∞ | 84.45 | 0.00 |
| L | 8 | -2998.19 | 6012.68 | 86.25 | 0.00 | 1.00 | ∞ | | |

Coefficients of the above analyses (naturally) averaged across the reduced confidence model set, including the conditional standard errors (SE) and 95% confidence limits (CL), the relative variable importance (*RI*) and number of models including each term (*N*).

| | Full confidence set | | | | | Excluding nested models | | | | |
|---|---|---|---|---|---|---|---|---|---|---|
| **Parameter** | **Estimate** | **SE** | **95% CL** | ***RI*** | ***N*** | **Estimate** | **SE** | **95% CL** | ***RI*** | ***N*** |
| (Intercept) | 1.33 | 0.14 | (1.06, 1.60) | | 6 | 1.33 | 0.14 | (1.05, 1.61) | | 6 |
| **N** | **0.69** | **0.09** | **(0.51, 0.86)** | **1.00** | **6** | **0.68** | **0.09** | **(0.51, 0.85)** | **1.00** | **4** |
| **E** | **0.43** | **0.09** | **(0.27, 0.60)** | **1.00** | **6** | **0.44** | **0.09** | **(0.27, 0.60)** | **1.00** | **4** |
| **N×E** | **0.48** | **0.16** | **(0.16, 0.79)** | **1.00** | **6** | **0.49** | **0.16** | **(0.18, 0.80)** | **0.95** | **1** |
| L | 0.16 | 0.16 | (-0.15, 0.46) | 0.60 | 5 | 0.14 | 0.16 | (-0.16, 0.45) | 0.03 | 1 |
| L×N | 0.25 | 0.17 | (-0.08, 0.58) | 0.33 | 3 | **0.33** | **0.17** | **(0.02, 0.66)** | **0.03** | **1** |
| L×E | 0.03 | 0.15 | (-0.27, 0.34) | 0.19 | 3 | | | | | |
| L×N×E | 0.23 | 0.32 | (-0.40, 0.85) | 0.04 | 1 | | | | | |

**Supplementary Table S12:** Results of the information-theoretic analyses examining the effects of the difference in sperm length (L), the difference in sperm number (N), the time to female sperm ejection (E), female SR length (S) and female thorax length (T) on $S_2$ within the SR ($N = 508$). Each model was a GLMM with the temporal block as a four-level random factor. An observation-level random effect was included to account for overdispersion. Due to many similar models, table only summarizes the data after excluding the slightly more complex, nested versions of higher-ranked models (1, 2). The most parsimonious confidence model set is highlighted in bold.

| Model | *df* | logLik | $AIC_c$ | $\Delta AIC_c$ | $w_i$ | cum $w_i$ | ER |
|---|---|---|---|---|---|---|---|
| **L + N + E + S + T + L×S + L×T + N×E** | **15** | **-2458.91** | **4948.80** | **0.00** | **0.25** | **0.25** | |
| **L + N + E + S + T + L×S + L×T** | **14** | **-2459.98** | **4948.80** | **0.01** | **0.25** | **0.51** | **1.00** |
| **L + N + E + S + T + L×N + L×T + N×E + N×T + L×N×T** | **17** | **-2457.23** | **4949.70** | **0.91** | **0.16** | **0.67** | **1.58** |
| **L + N + E + S + T + L×T + N×E** | **14** | **-2460.93** | **4950.70** | **1.92** | **0.10** | **0.76** | **2.61** |
| **L + N + E + S + T + L×T** | **13** | **-2462.05** | **4950.80** | **2.04** | **0.09** | **0.85** | **2.78** |
| **N + E + S + T + N×E** | **12** | **-2463.35** | **4951.30** | **2.53** | **0.07** | **0.92** | **3.56** |
| **N + E + S + T** | **11** | **-2464.58** | **4951.70** | **2.88** | **0.06** | **0.98** | **4.22** |
| N + E + S + N×E | 11 | -2466.78 | 4956.10 | 7.29 | 0.01 | 0.99 | 36.14 |
| N + E + S | 10 | -2467.98 | 4956.40 | 7.60 | 0.01 | 1.00 | 42.17 |
| L + N + E + T + L×N + L×T + N×E + N×T + L×N×T | 16 | -2463.17 | 4959.40 | 10.64 | 0.00 | 1.00 | ∞ |
| N + E + N×E | 10 | -2469.53 | 4959.50 | 10.71 | 0.00 | 1.00 | ∞ |
| N + E | 9 | -2470.77 | 4959.90 | 11.09 | 0.00 | 1.00 | ∞ |
| L + E + S + T + L×S + L×T | 13 | -2477.70 | 4982.10 | 33.33 | 0.00 | 1.00 | ∞ |
| E + S + T | 10 | -2482.03 | 4984.50 | 35.70 | 0.00 | 1.00 | ∞ |
| E + S + E×S | 10 | -2484.91 | 4990.30 | 41.47 | 0.00 | 1.00 | ∞ |
| E + S | 9 | -2486.07 | 4990.50 | 41.70 | 0.00 | 1.00 | ∞ |
| E | 8 | -2489.05 | 4994.40 | 45.60 | 0.00 | 1.00 | ∞ |
| L + N + S + T + L×N + L×S + L×T + N×T + L×N×T | 16 | -2480.76 | 4994.60 | 45.84 | 0.00 | 1.00 | ∞ |
| N + S | 9 | -2488.16 | 4994.70 | 45.89 | 0.00 | 1.00 | ∞ |
| N | 8 | -2492.40 | 5001.10 | 52.29 | 0.00 | 1.00 | ∞ |
| S | 8 | -2503.59 | 5023.50 | 74.66 | 0.00 | 1.00 | ∞ |
| (Null) | 7 | -2508.79 | 5031.80 | 83.01 | 0.00 | 1.00 | ∞ |

Coefficients of the above analyses (naturally) averaged across the reduced confidence model set, including the conditional standard errors (SE) and 95% confidence limits (CL), the relative variable importance (*RI*) and number of models including each term (*N*).

| Parameter | Estimate | SE | 95% CL | *RI* | *N* |
|---|---|---|---|---|---|
| (Intercept) | 2.35 | 0.17 | (2.02, 2.68) | | 7 |
| **N** | **0.76** | **0.12** | **(0.51, 1.00)** | **1.00** | **5** |
| **E** | **0.87** | **0.12** | **(0.63, 1.11)** | **1.00** | **7** |
| **S** | **-0.56** | **0.12** | **(-0.79, -0.32)** | **1.00** | **7** |
| **T** | **0.35** | **0.12** | **(0.11, 0.59)** | **1.00** | **7** |
| **L×T** | **-0.55** | **0.24** | **(-1.02, -0.09)** | **0.87** | **2** |
| L | -0.10 | 0.17 | (-0.43, 0.22) | 0.87 | 7 |
| N×E | 0.40 | 0.25 | (-0.09, 0.88) | 0.59 | 5 |
| **L×S** | **0.48** | **0.24** | **(0.02, 0.94)** | **0.51** | **4** |
| **L×N×T** | **-1.12** | **0.47** | **(-2.03, -0.21)** | **0.16** | **1** |
| N×T | 0.21 | 0.24 | (-0.26, 0.69) | 0.16 | 1 |
| L×N | 0.18 | 0.25 | (-0.31, 0.66) | 0.16 | 1 |

References

1. K. P. Burnham, D. R. Anderson, *Model Selection and Multi-Model Inference: A Practical Information-Theoretic Approach* (Springer, New York, 2002).

2. S. A. Richards, M. J. Whittingham, P. A. Stephens, Model selection and model averaging in behavioural ecology: The utility of the IT-AIC framework. *Behav. Ecol. Sociobiol.* **65**, 77–89 (2011).
